# Loss of Mitochondrial Enoyl CoA Reductase causes elevated ceramide levels and impairs iron metabolism

**DOI:** 10.1101/2023.04.14.536458

**Authors:** Debdeep Dutta, Oguz Kanca, Seul Kee Byeon, Paul C. Marcogliese, Zhongyuan Zuo, Rishi V. Shridharan, Jun Hyoung Park, Undiagnosed Diseases Network, Guang Lin, Ming Ge, Gali Heimer, Jennefer N. Kohler, Matthew T. Wheeler, Benny A. Kaipparettu, Akhilesh Pandey, Hugo J. Bellen

## Abstract

In most eukaryotic cells fatty acid synthesis occurs in the cytoplasm as well as in mitochondria. However, the relative contribution of mitochondrial fatty acid synthesis (mtFAS) to the cellular lipidome of metazoans is ill-defined. Hence, we studied the function of the fly Mitochondria enoyl CoA reductase (Mecr), the enzyme required for the last step of mtFAS. Loss of *mecr* causes lethality while neuronal loss leads to progressive neurological defects. We observe an elevated level of ceramides, a defect in Fe-S cluster biogenesis and increased iron levels in *mecr* mutants. Reducing the levels of either iron or ceramide suppresses the neurodegenerative phenotypes indicating that increased ceramides and iron metabolism are interrelated and play an important role in the pathogenesis. Mutations in human *MECR* cause pediatric-onset neurodegeneration and patient-derived fibroblasts display similar elevated ceramide levels and impaired iron homeostasis. In summary, this study shows an as-yet-unidentified role of *mecr/MECR* in ceramide and iron metabolism providing a mechanistic link between mtFAS and neurodegeneration.

## Introduction

The synthesis of fatty acids in eukaryotic cells relies on two independent pathways. The first operates in the cytoplasm (FAS I) and relies on Fatty Acid Synthase, an enzyme that carries six enzymatic domains for fatty acid synthesis ^1,2^. The second pathway operates in mitochondria (FAS II/mtFAS) and involves six different enzymes. *MECR* (***M****itochondrial **E**noyl **C**oA **R**eductase*) encodes one of these enzymes which is required for the final step of mitochondrial fatty acid synthesis in humans (Extended Data Fig. 1a).

Since the discovery of mtFAS, its role in cellular growth and differentiation has been documented ^3–8^. In yeast, loss of *ETR*, the homolog of human *MECR*, leads to mitochondrial dysfunction and respiratory growth arrest ^7, 8^. In mice, loss of *Mecr* is embryonic lethal. However, a Purkinje cell-specific conditional knock-out leads to ataxia in nine months old mice ^4, 9^. In murine C2C12 myoblasts, reduced mtFAS activity impairs their differentiation ^6^. Recently, bi-allelic mutations in *MECR* have been shown to cause a pediatric-onset neurodegenerative disorder called MEPAN (**M**itochondrial **E**noyl Reductase **P**rotein **A**ssociated **N**eurodegeneration; OMIM # 617282) syndrome ^10–12^. Patients with MEPAN present in early childhood (1-6.5 years) with basal ganglia lesions leading to dystonia, chorea and other movement disorders, followed by progressive optic atrophy and occasional developmental delay. Finally, mtFAS dysfunction has also been implicated in Parkinson’s disease ^13, 14^. Neurological symptoms in MEPAN indicates the importance of mtFAS in neuronal survival and maintenance, and raise an important question: how does mtFAS contribute to neuronal maintenance?

The cytoplasmic fatty acid synthesis pathway produces palmitate (C16) linked to the CoA cofactor, which is used for the synthesis of lipids including phospholipids and sphingolipids. In contrast, mtFAS produces acyl chains of varying carbon length (C4 to C18) linked to mitochondrial Acyl Carrier Protein (mtACP/ACP) ^15–18^. Lipoic acid, an essential cofactor for multiple mitochondrial enzymes ^19^ is derived from octanoyl (C8)-ACP ^17, 20^. However, the role of the other Acyl-ACPs in lipid synthesis is not well defined. Indeed, the consequences of impaired mtFAS on the cellular lipidome have remained elusive based on studies in yeast, Neurospora, and mammalian cell lines ^6, 21–23^. For example, loss of *Mecr* in murine C2C12 myoblast cells was reported to not affect any major cellular phospholipids ^6^. However, a reduction of mtACP caused decreased levels of lysophospholipids and sphingolipids in HeLa cells ^23^. In contrast, in *Neurospora crassa*, mtACP mutants displayed an increase in mitochondrial lysophospholipid levels ^21^. In *Trypansosoma bruci*, RNAi mediated knockdown of mtACP decreased the phosphoinositides and phosphoethanolamine levels but it did not alter sphingomyelin levels ^22^. Finally, upon loss of mtACP in yeast, no major difference in mitochondrial lipids and fatty acids was noted ^21^. Given that many enzymes and enzymatic products are involved in mtFAS, loss of any of the six genes in the pathway may cause different phenotypic signatures. Additionally, the phenotypic differences may arise because the above-mentioned studies were carried out using different model systems in different culture conditions, at different time points and with alleles of different strength.

To explore the molecular consequences of impaired mtFAS, we studied the effects of *mecr*/*MECR* loss in fruit flies and fibroblasts from MEPAN patients. Loss of *mecr* in flies causes larval lethality and the neuron specific loss/reduction of *mecr* progressively impairs locomotion, synaptic transmission, and phototransduction in adult flies. Surprisingly, *mecr* loss results in elevated ceramide levels in flies. Similarly, fibroblasts derived from individuals with MEPAN also display elevated levels of ceramide. This increase in ceramide in flies and in human cells is associated with a defect in iron-sulfur (Fe-S) cluster synthesis and iron metabolism. Importantly, these defects are rescued by lowering the levels of either ceramide or iron indicating an interrelationship between the two metabolic pathways. In summary, our study identifies an evolutionarily conserved role of Mecr/MECR/mtFAS in ceramide and iron metabolism that underlies the demise of neurons.

## Results

### Human *MECR* can functionally replace fly *mecr*

Since human MECR is required for the last step of mtFAS (Extended Data Fig. 1a), we evaluated the consequences of the loss of mtFAS in fruit flies by targeting CG16935/*mecr*, an as-yet uncharacterized gene in flies that encodes the Mecr protein. We used CRISPR mediated homologous recombination to generate a strong loss-of-function allele (*mecr^TG4^*). An artificial exon encoding a *Splice-Acceptor-T2A-GAL4-polyA* construct was introduced in the coding intron of the gene (Extended Data Fig. 1b) ^24^. The *polyA* signal arrests transcription causing the endogenous fly mRNA to be truncated. Hence, *T2A-GAL4* alleles are typically strong loss-of-function mutations ^25^. Indeed, in our Real-time PCR data using two independent primer sets, we detected no *mecr* transcript in the *mecr^TG4^* mutants (Extended Data Fig. 1c). We also used another insertion allele, *mecr^A^*, in which a PiggyBac transposon, which contains an artificial exon that introduces a STOP codon is inserted in the coding intron (Extended Data Fig. 1b) ^26^. *mecr^A^* and *mecr^TG4^* alleles are homozygous and transheterozygous lethal. Moreover, they are lethal over a molecularly defined deficiency (*Df(2R)BSC307/CyO*) that removes the *mecr* coding region (Fig. 1a) indicating that the observed lethality is due to loss of *mecr* function. All homozygous and transheterozygous animals die as late 2nd instar/early 3rd instar larvae suggesting that *mecr^TG4^* and *mecr^A^*are strong loss-of-function alleles. A genomic duplication construct that harbors the *mecr* coding and regulatory region (Genomic rescue construct, *GR*) (P[acman] *CH321-09M04*) ^27^ as well as expression of the fly cDNA using the ubiquitous *daughterless (da)-GAL4* fully rescue the lethality (Fig. 1a and Extended Data Fig. 1d), showing that the phenotypes associated with *mecr^TG4^* are due to loss of *mecr* function. The fly Mecr protein is 44% identical to and shares 59% similarity with the human MECR protein with a 15/15 DIOPT score ^28, 29^. Moreover, all the amino acids that have missense variants in MEPAN patients are conserved in the fly protein (Extended Data Fig. 2a). Importantly, the reference human *MECR* cDNA rescues the lethality of homozygous *mecr^TG4^*mutants showing functional conservation (Fig. 1a). The previously reported human *MECR* nonsense variant, MECR^Tyr285*^, failed to rescue lethality whereas two missense variants, MECR^Gly232Glu^ and MECR^Arg258Trp^ partially rescue lethality (Extended Data Fig. 2b). These and other data^10^ argue that MEPAN patients typically carry one strong and one weak loss-of-function alleles. In summary, *mecr* encodes an essential protein that is functionally conserved between flies and humans.

**Figure 1:**
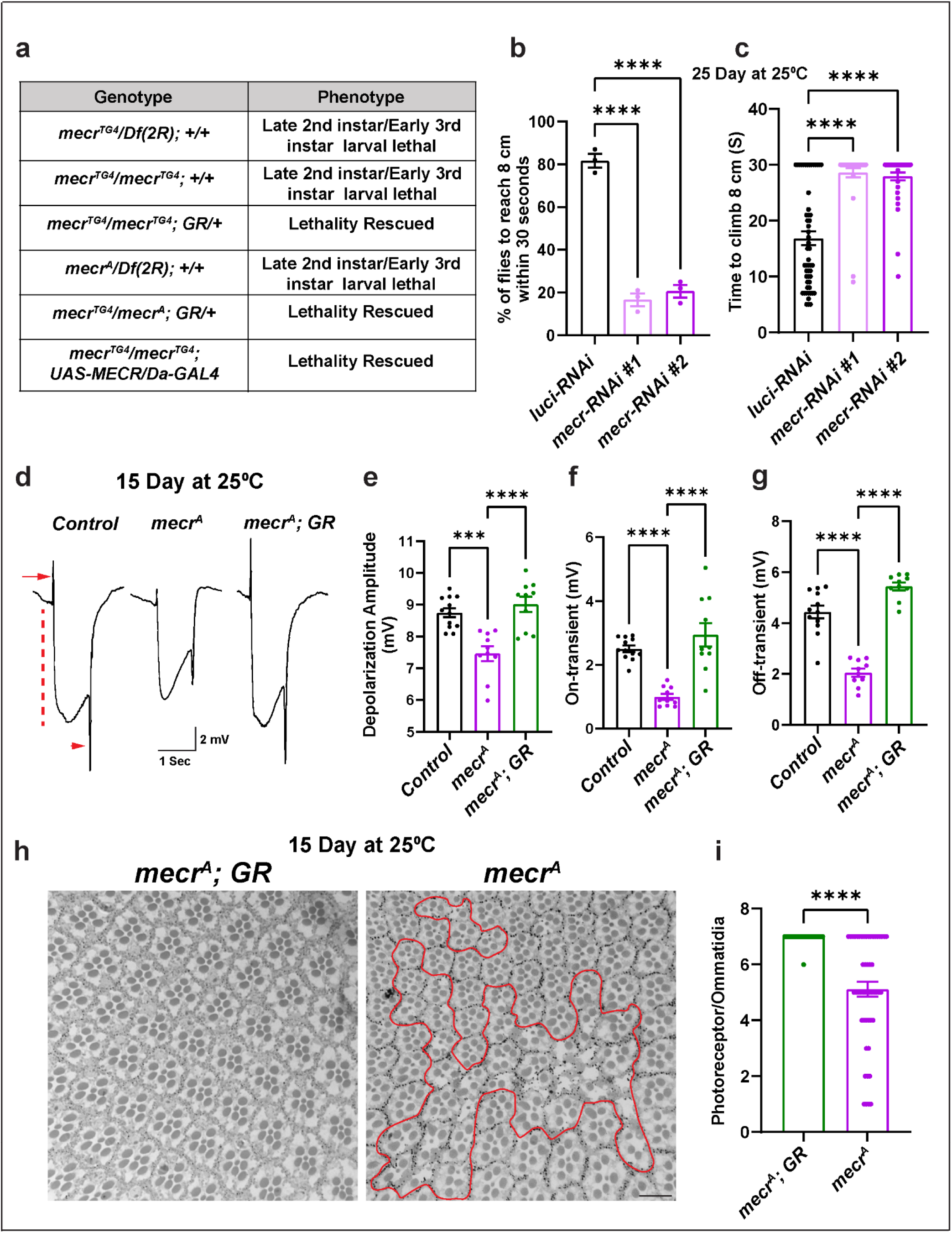
Loss of *mecr* causes an age-dependent locomotor impairment, ERG defects and photoreceptor loss. (a) The phenotypes in *mecr* mutants and complementation with an 80 kb P[acman] rescue construct (*GR*)^27^ containing the *mecr* gene or ubiquitous expression of human *MECR*. (b-c) Climbing ability of 25-day-old flies upon neuronal knockdown (by *elav-GAL4*) of *mecr* with two independent RNAi lines. (b) Percentage of 25-day-old flies that are able to climb 8 cm within 30 seconds. (c) Average time taken by 25-day-old flies to climb 8 cm. Total number of flies counted for three independent replicates were: n = 54 (*luci-RNAi*), n= 35 (*mecr-RNAi#1*), n=46 (*mecr-RNAi#2*). (d) ERG traces of controls and *mecr^A^* mutant clones of 15-day-old flies. ERG recordings are induced by light and exhibit “on” and “off” transients (arrow and arrowhead), indicative of synaptic communication between the PR neurons and postsynaptic cells. They also exhibit a corneal negative response, the amplitude of which corresponds to the depolarization of PR neurons (dashed line). (e-g) Quantification of depolarization amplitude as well as “on” and “off” transients for the respective genotypes. n = 12 (*Control*), n=10 (*mecr^A^*), n=10 (*mecr^A^; GR*). (h) Retinal sections and (i) quantification of PRs per ommatidium from control and *mecr^A^* mutant clones in 15-day-old flies. n=51 (*mecr^A^* and *mecr^A^; GR*). The red line marks the clonal area. All the flies were maintained at 25°C. For statistical analyses between two samples, two-tailed Student’s t test, and for three samples, one-way ANOVA followed by a Tukey’s post-hoc test are carried out. Error bars represent SEM (***p < 0.001; ****p < 0.0001).

## Neuronal knockdown of *mecr* induces an age-dependent climbing defects in flies

Given the lethality associated with the loss of *mecr*, we evaluated the effects of neuronal knockdown of *mecr* on locomotor ability of aged flies. We used the pan-neuronal driver, *elav-GAL4* to perform RNAi-mediated knockdown of *mecr* in neurons. We used two independent RNAi lines which reduce the levels of Mecr protein by ∼45% (RNAi#1) and ∼75% (RNAi#2) (Extended Data Fig. 3a-b) at 25°C. Hence, we used RNAi#2 for the remaining experiments. Although neuronal knock-down of *mecr* does not cause obvious climbing defects in young flies (3 days post-eclosion), a significant impairment in climbing was observed in aged flies (25-days post-eclosion) (Fig. 1b-c). Compared to control flies, where ∼ 80% flies climb to 8 cm within 30 seconds, only ∼20% of flies with neuronal *mecr* knock-down were able to climb at day 25 (Fig. 1b). The average climbing time was also significantly increased in these flies compared to age-matched control flies (Fig. 1c). In addition, we noted a mild but significant lifespan reduction in these flies with neuronal *mecr* knock-down (Extended Data Fig. 3c). These results indicate an age-dependent loss of neuronal function when *mecr* levels are reduced. In summary, neuronal knockdown of *mecr* causes a progressive loss of motor function.

## Loss of *mecr* causes the demise of photoreceptor neurons

To evaluate the impact of *mecr* loss on phototransduction and synaptic transmission, we performed electroretinograms (ERG). Given that *mecr* mutants are homozygous lethal, we generated mutant clones in the eye using the *Flp-FRT* system (*FRT42B mecr^A^ / FRT42B Ubi-GFPnls; ey-Flp/+*) ^30,31^ and performed ERGs by placing the electrode in the mutant eye clones. In young flies (3-5 days old), we did not observe any difference in ERG traces between mutant and control tissue (Extended Data Fig. 3d-f). However, 15-day-old flies exhibit decreased phototransduction amplitudes indicating a reduced ability of photoreceptors (PRs) to sense light (Fig. 1d-e). Moreover, the ERG traces also exhibit reduced ON and OFF transients, suggesting a defect in synaptic communication between the photoreceptors and the postsynaptic neurons (Fig. 1f-g). To determine if PR morphology is affected in *mecr* mutant clones, we imaged 15-day old retinas. As shown in Fig. 1h-i, many ommatidia exhibit an aberrant morphology with fewer PR neurons in the mutant clones. Hence, there is a progressive loss of neuronal function and photoreceptors upon loss of *mecr*.

## *mecr* encodes a mitochondria-localized protein in fruit fly

The subcellular localization of Mecr protein *in vivo* has not been determined previously in multicellular organisms. In yeast, the Etr protein was reported to be predominantly localized in the nucleus ^32^. However, upon overexpression of the gene, the protein was also shown to localize to mitochondria ^8^. Other overexpression studies in HeLa cells, indicated that Mecr was localized to mitochondria, cytoplasm, as well as nuclei ^3, 33–35^. To determine the endogenous subcellular localization of Mecr protein in fruit fly, we used a genomic rescue construct of *mecr* that is tagged with a C-terminal GFP (CBGtg9060C0290D) ^36^ (Fig. 2a). Importantly, this tag does not disrupt the function of *mecr* as it rescues the lethality of *mecr* mutants, and the rescued flies appear healthy with no obvious phenotypes (Extended Data Fig. 1d). Interestingly, we observe a higher expression level of this protein in some glial subsets when compared to neurons in the larval brain (Fig. 2b). Staining with antibodies against GFP and ATP5α, an inner mitochondrial membrane protein ^37, 38^ shows that Mecr localizes to mitochondria in multiple larval tissues (Fig. 2c & Extended Data Fig. 3g). To determine the localization of human MECR protein, we expressed the human gene with the *actin-GAL4* driver in S2 cells, stained with an antibody against MECR (AB_1853714), and observed mitochondrial co-localization of MECR and ATP5α (Extended Data Fig. 3h). In summary, fly and human Mecr/MECR proteins are localized to mitochondria.

**Figure 2:**
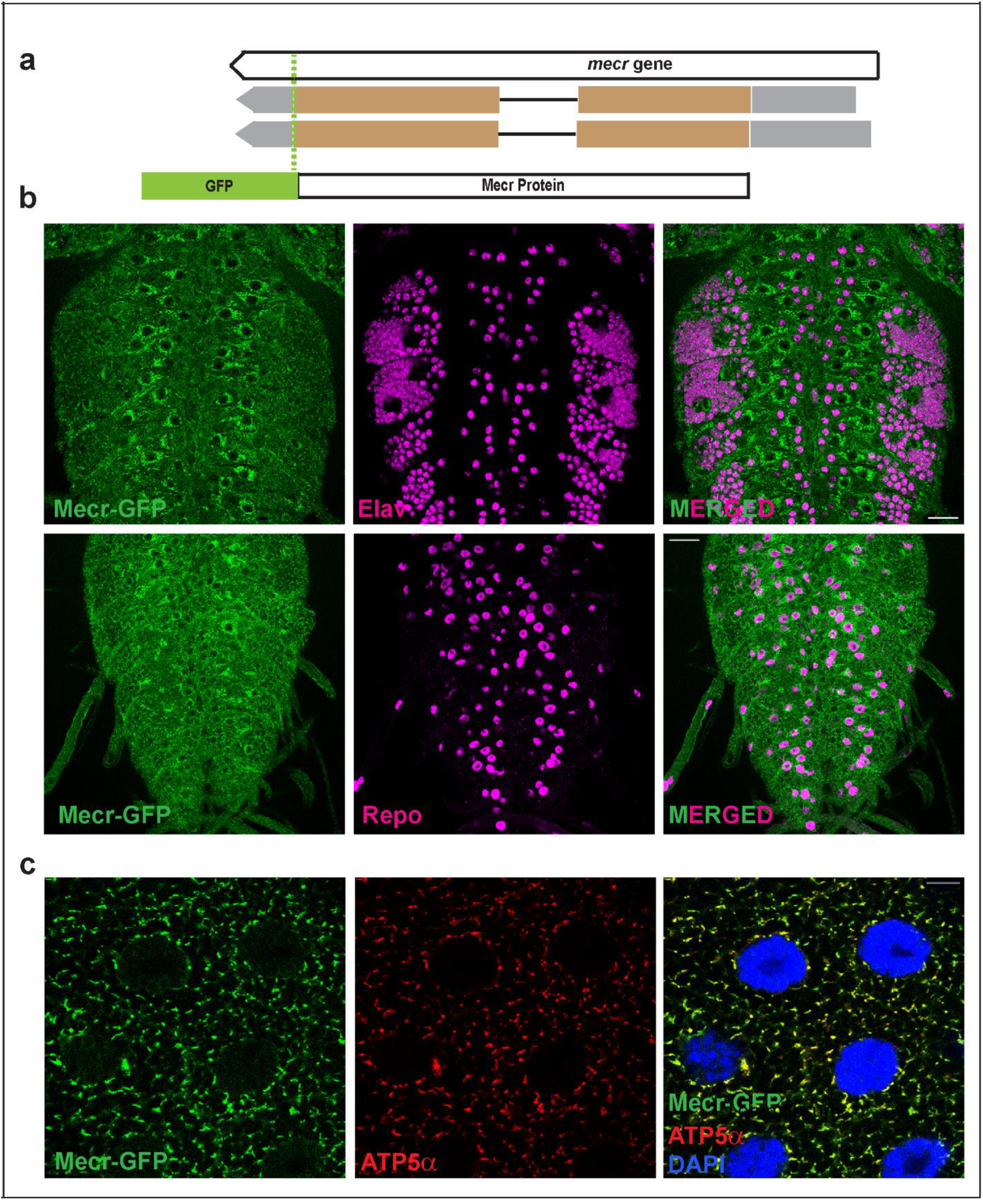
Mecr is enriched in a subset of glial cells in larval brain and in salivary gland. (a) Schematic diagram showing the *mecr-GFP* allele (Fosmid clone, CBGtg9060C0290D)^36^ used in this study. (b) Expression of Mecr-GFP in Elav-positive neuronal and Repo-positive glial cells of the 3^rd^ instar larval brain. Scale bar: 25 µm. (c) Salivary gland cells from 3rd instar larva and colocalization of ATP5α, an inner mitochondrial membrane protein, and Mecr-GFP. Scale bar: 10 µm.

## Loss of Mecr/ MECR leads to ceramide accumulation

Given that loss of MECR affects mtFAS, we assumed that loss of *mecr* or *MECR* would reduce at least some of the lipids of the cell. We performed a comparative lipidomic analysis using *mecr^TG4^* homozygous mutant larvae and human patient-derived fibroblasts. We used fibroblasts derived from two MEPAN patients (Patient 1 and Patient 2, Siblings) and their healthy parent (Extended Data Fig. 4a) identified through the Undiagnosed Diseases Network ^39^. Both patients carry variants that lead to a significant reduction in *MECR* transcript levels (Extended Data Fig. 4b) and have developmental delay, movement disorders and hypotonia. An untargeted lipidomic approach was taken to measure the major phospholipid and sphingolipid species. In *mecr* fly mutants, we observed increased levels of some phospholipids including phosphatidylethanolamine (PE), phosphatidylinositol (PI) and phosphatidylglycerol (PG) but no significant change in phosphatidylcholine (PC) and phosphatidylserine (PS) levels (Extended Data Fig. 4h-l). However, we did not observe any significant alterations in the levels of any major phospholipid species including PC, PE, PI, PS and PG in patient-derived fibroblasts (Extended Data Fig. 4c-g) consistent with the murine C2C12 cells with a known defect in mtFAS ^6^. In contrast, the levels of ceramides, a subclass of sphingolipids, were significantly elevated in the *mecr* mutants as well as in MEPAN patient-derived fibroblasts (Fig. 3a and d). In summary, none of the lipids based on lipidomics assays are decreased. Rather, we observe a systematic increase in ceramides.

**Figure 3:**
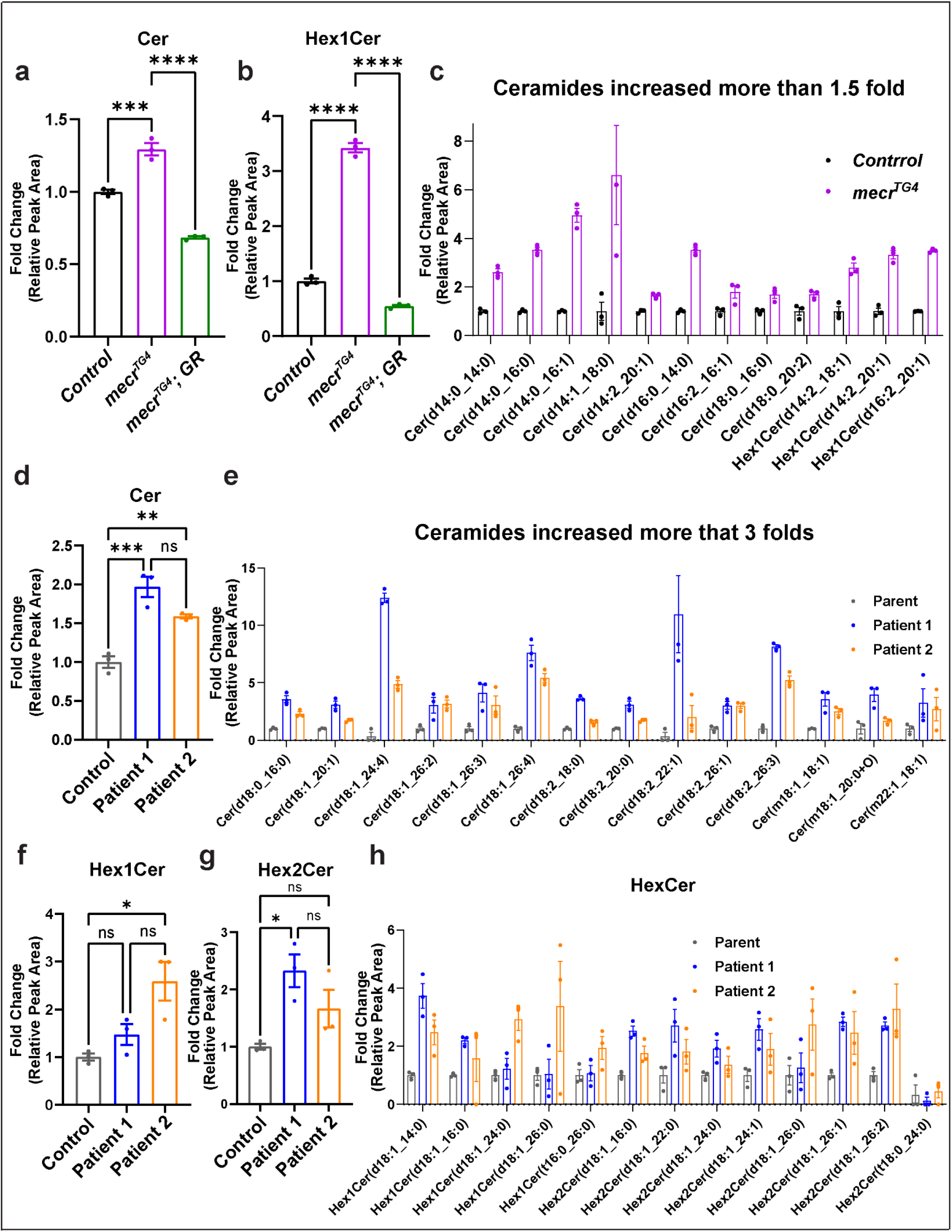
Loss of *mecr/MECR* leads to an increase in ceramide levels. (a-c) Relative amounts of ceramides and hexosylceramides in *mecr^TG4^*mutants. (d-e) Relative amounts of ceramides in fibroblasts from patients. (f-h) Relative amount of hexosylceramides in patient-derived fibroblasts. For statistical analyses, one-way ANOVA followed by a Tukey’s post-hoc test is carried out. Error bars represent SEM (*p < 0.05; **p < 0.01; ***p < 0.001; ****p < 0.0001).

Ceramides are a central hub for sphingolipid metabolism and are responsible for the production of glucosylceramide as well as galactosylceramide, collectively referred to as hexosylceramides (HexCer) ^40, 41^. The levels of ceramides (including HexCer species) were significantly increased in the *mecr^TG4^* mutants compared to animals carrying a genomic rescue construct or *yw* controls (Fig. 3a-b). 25 out of 33 ceramide species analyzed, including HexCer, were increased in fly mutants (Fig. 3c and Extended Data Fig. 5a). We next explored the ceramide levels in patient-derived fibroblasts. The total levels of ceramides are increased about two-fold when compared to control fibroblasts (Fig. 3d). Out of 46 ceramide species analyzed, 43 are increased in fibroblasts from both patients (Fig. 3e and Extended Data Fig. 5b). Interestingly, some of the ceramide species (d18:1_24:4, d18:1_26:4, d18:2_22:1, and d18:2_26:3) are increased ten-fold or more in patient 1. The levels of Hex1Cer (a Ceramide with one sugar) and Hex2Cer (a Ceramide with two sugars) are also elevated in cells from both patients. However, Hex1Cer reaches a statistical significance level in patient 1 while Hex2Cer reaches a statistical significance level in patient 2 (Fig. 3f-g). Indeed, 12 out of 13 HexCer species analyzed were increased in the patient fibroblasts when compared to the parent control (Fig. 3h). In summary, we observe a significant increase of ceramides upon loss of *mecr*/*MECR* in fly mutants and human patient cells, suggesting a conserved role for this protein in ceramide metabolism.

### Reducing ceramide levels alleviate neurodegeneration in adult flies

The availability of drugs that reduce ceramide levels provides an opportunity to lower ceramides and assess their effect on the neurodegenerative phenotypes *in vivo*. We first assessed ceramide levels in 25-day-old adult fly heads upon neuronal knockdown of *mecr* by using an antibody against ceramides (MID 15B4)^42^. We noticed an increased level of ceramides in the heads of 25-day-old flies (Fig. 4a). As shown in Fig. 1b-c and 4b-c, these flies show obvious climbing defects. We treated these flies with myriocin, to reduce the enzymatic activity of Serine Palmitoyl Transferase (SPT), which acts in the rate-limiting step of the *de novo* ceramide synthesis ^43^. We also treated these *mecr* knockdown flies with desipramine, a drug that inhibits the lysosomal sphingomyelinases and reduces the production of ceramides through the salvage pathway in lysosomes ^44^. Upon treatment with either of these drugs, the climbing ability was significantly improved in 25-day-old flies (>70% flies) (Fig. 4b-c). We observed a similar improvement in ERGs in 25-day-old flies. Treatment with either of these drugs also alleviates the synaptic transmission defects upon neuronal knockdown of *mecr* (Fig. 4d-f). In summary, decreasing ceramide levels is beneficial for neuronal function when *mecr* levels are reduced.

**Figure 4:**
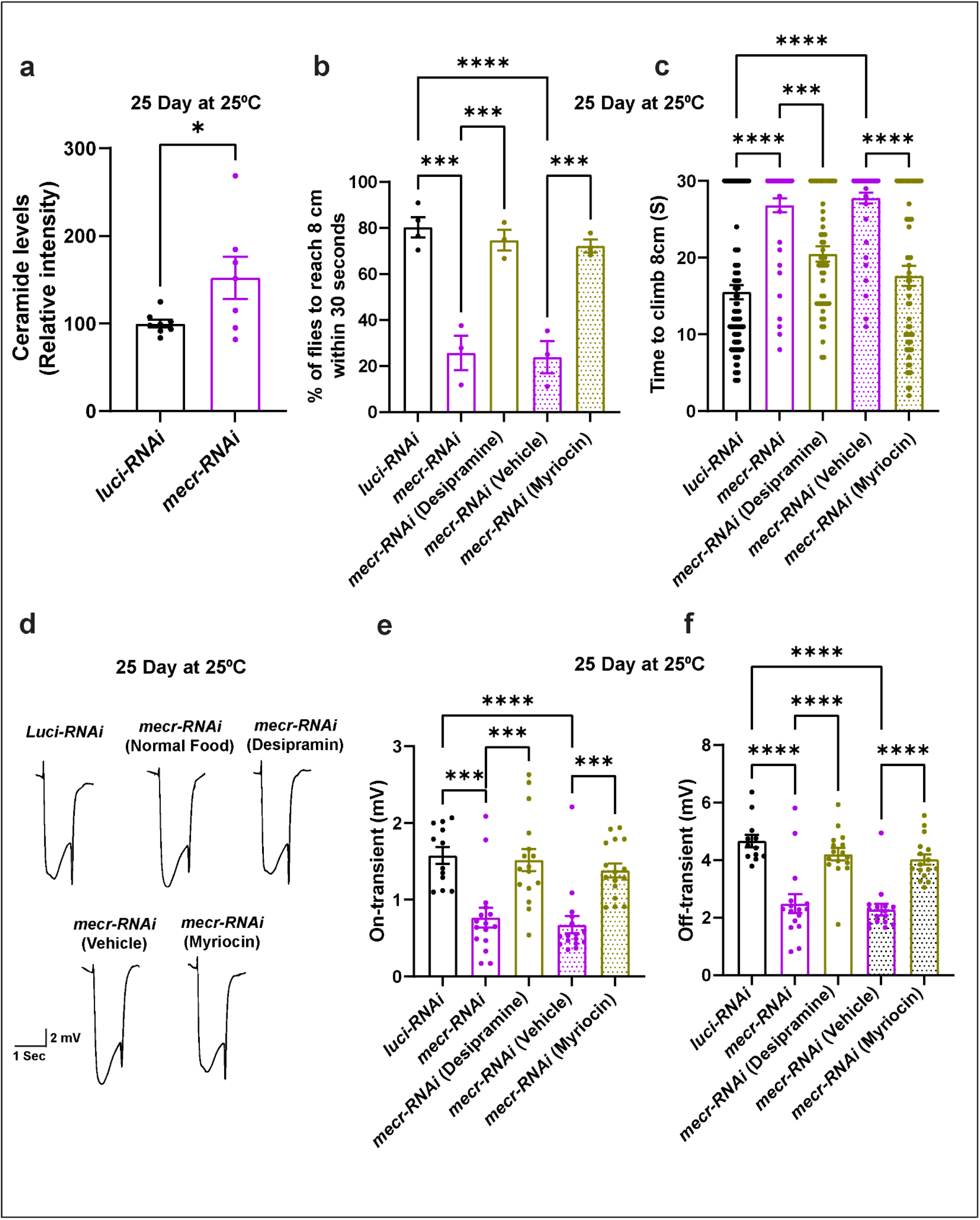
Reducing Ceramide levels alleviates the age dependent phenotypes in fruit fly. (a) Quantification of relative ceramide intensity in 25-day-old brains upon neuronal knockdown (by *elav-GAL4*) of *mecr*. n = 8 (*luci-RNAi*), n = 7 (*mecr-RNAi*) biologically independent samples. (b-c) Average percentage and climbing time of flies with and without Myriocine and Desipramine treatment. Total number of flies counted for three independent replicates were: n = 84 (*luci-RNAi*), n = 51 (*mecr-RNAi*), n = 51 (*mecr-RNAi* with Desipramine treatment), n = 51 (*mecr-RNAi* with vehicle) and n = 57 (*mecr-RNAi* with Myriocin treatment). (d-f) ERG traces and quantification showing the effects of Myriocine and Desipramine treatment on 25-day-old flies with neuronal knockdown of *mecr*. n = 12 (*luci-RNAi*), n = 16 (*mecr-RNAi*), n = 16 (*mecr-RNAi* with Desipramine treatment), n = 16 (*mecr-RNAi* with vehicle) and n = 16 (*mecr-RNAi* with Myriocin treatment). For statistical analyses between two samples, two-tailed Student’s t test, and for three samples, one-way ANOVA followed by a Tukey’s post-hoc test are carried out. Error bars represent SEM (*p < 0.05; ***p < 0.001; ****p < 0.0001).

### Loss of *mecr/MECR* affects mitochondrial function and morphology in flies and human fibroblasts

To evaluate the mitochondrial function in *mecr* mutant larvae, we measured ATP production ^45–47^. Loss of *mecr* reduces ATP production by ∼50% when compared to controls (Fig. 5a). One copy of the genomic rescue construct in the homozygous mutant background improved ATP production (Fig. 5a). Given that impaired ATP production can be associated with loss of mitochondrial membrane potential ^48^, we assessed mitochondrial membrane potential in larval muscles by using tetramethylrhodamine ethyl ester (TMRE) ^49^. TMRE is incorporated between the inner and outer mitochondrial membrane in healthy mitochondria, but when the mitochondrial membrane potential is reduced, TMRE does not intercalate efficiently, leading to a reduction in the intensity of staining. Homozygous *mecr^TG4/TG4^* larval mutant muscles show a significant decrease in mitochondrial membrane potential (Fig. 5b-c). Finally, we assessed the activity of the various electron transport chain (ETC) enzyme complexes in homozygous *mecr^TG4^* second instar larvae and observed reduced activity of Complex-I, I+III and IV and increased activity of Complex-II (Extended Data Fig. 6a). Additionally, we assessed the mitochondrial morphology in photoreceptor neurons of *mecr* clones through transmission electron microscopy (TEM). The mitochondria in these neurons are less electron-dense and display a severe disruption of cristae (Fig. 5d). In summary, these data show a severe dysfunction of mitochondria associated with loss of *mecr* and morphological impairment of neuronal mitochondria when *mecr* is lost in eye clones.

**Figure 5:**
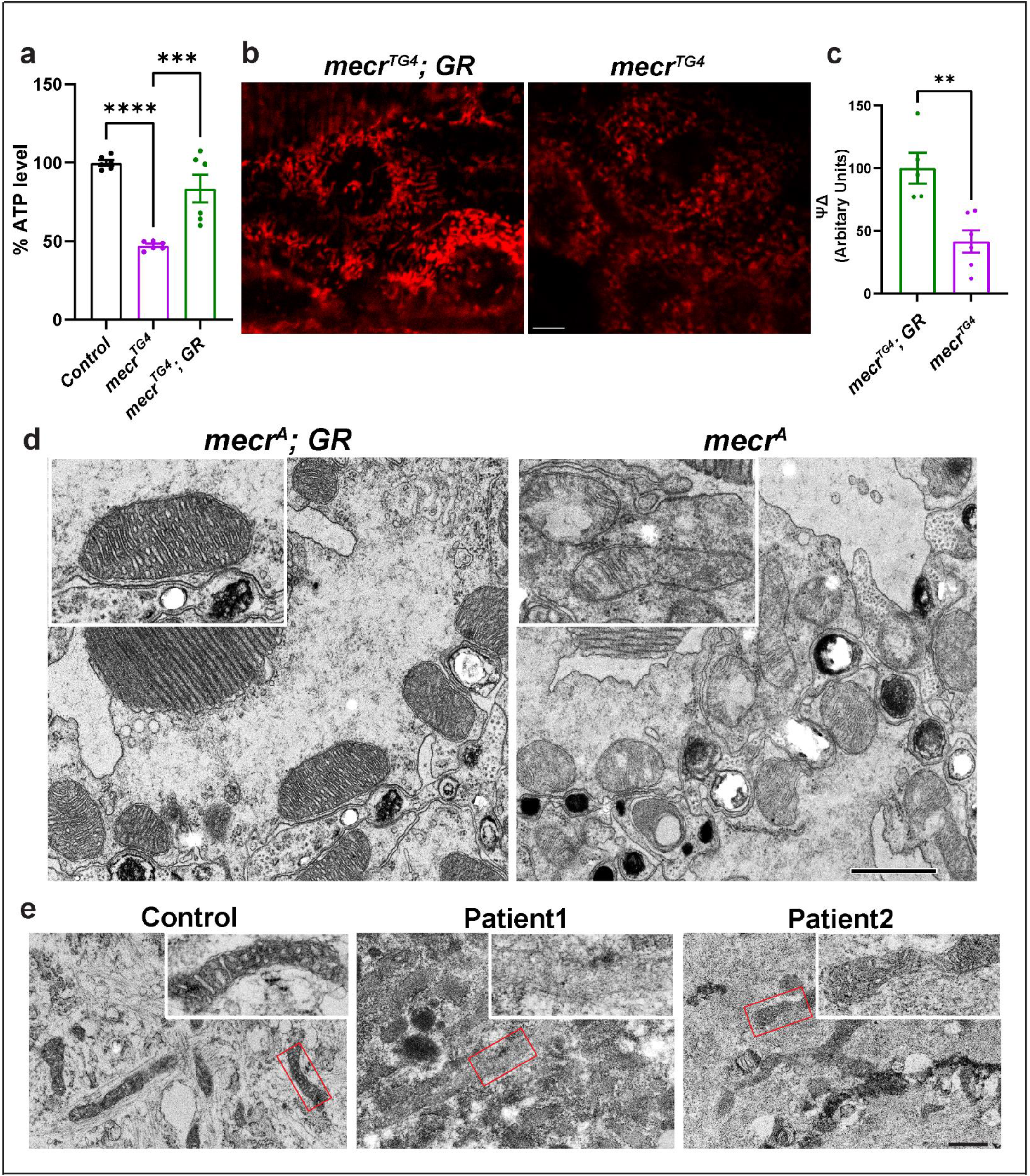
Loss of *mecr/MECR* impairs mitochondrial function and morphology. (a) Relative levels of ATP in *mecr^TG^*^4^ mutants and controls. (b-c) Relative levels of mitochondrial membrane potential as measured by TMRM in larval muscle. Scale bar is 5 µm. n = 5 (*mecr^TG4^; GR*), n = 6 (*mecr^TG4^*) biologically independent samples. (d) Transmission Electron micrographs showing mitochondrial morphology in photoreceptor neurons of 15-day-old flies. Inset: Single mitochondrion showing the extent of severity in mitochondrial morphology in *mecr* mutant clones (right) compared to control (left). Scale bar 0.8 µm. (e) Transmission Electron micrographs showing mitochondrial morphology in control and MEPAN patient derived fibroblasts. Inset: Enlarged area from the indicated regions shown in e. Scale bar 0.6 µm. For statistical analyses between two samples, two-tailed Student’s t test, and for three samples, one-way ANOVA followed by a Tukey’s post-hoc test are carried out. Error bars represent SEM (**p < 0.01; ***p < 0.001; ****p < 0.0001).

We then assessed the mitochondrial activity in fibroblasts from MEPAN patients. The ATP levels are decreased in the fibroblasts of the affected patients by ∼50% when compared to the parent control (Extended Data Fig. 6b). Previously, the oxygen consumption rate was measured in fibroblasts from three patients, and a mild and variable reduction in a subset of respiratory parameters was noted ^10^. We assessed basal respiration, maximal respiration and spare respiratory capacity using Seahorse assays ^50^ in fibroblasts from the patients and the parental control. As shown in Extended Data Fig. 6c-e, all were significantly reduced in both patients when compared to the parental fibroblasts. These data are consistent with Seahorse assays performed on mice C2C12 cells compromised in mtFAS ^6^. We also evaluated the mitochondrial morphology in patient-derived fibroblasts through transmission electron microscopy. The electron density of mitochondria is significantly reduced, and the morphology of mitochondria is impaired in fibroblasts from both patients (Fig. 5e). The fibroblasts from patient 1 are more severely affected than those of patient 2 (Fig. 5e). In summary, our data show that *mecr*/*MECR* is required for proper morphology and bioenergetic functions of mitochondria in fruit flies and human cells.

### Loss of Mecr impairs iron metabolism

Next, we attempted to define the molecular mechanism by which the loss of Mecr affects ceramide levels. An increase in ceramides levels can occur when: i) the *de novo* synthesis of ceramides is increased in the ER ^51, 52^, ii) the salvage pathway is activated in lysosomes^44^, or iii) iron metabolism is impaired in mitochondria ^53–55^. Previously, we discovered that loss-of-function mutation in Frataxin, a mitochondrial protein required for Fe-S cluster assembly ^56^, lead to increased iron levels and accumulation of ceramides ^53, 54^. This prompted us to test iron levels in the fly *mecr* mutants. A ferrozine-based colorimetric assay revealed an increased level of iron upon loss of *mecr* in flies when compared to controls (Fig. 6a). Additionally, Pearl’s staining ^53, 54, 57^ revealed an iron accumulation in the brain neuropils in two-day old mutant larvae (Fig. 6b).

**Figure 6:**
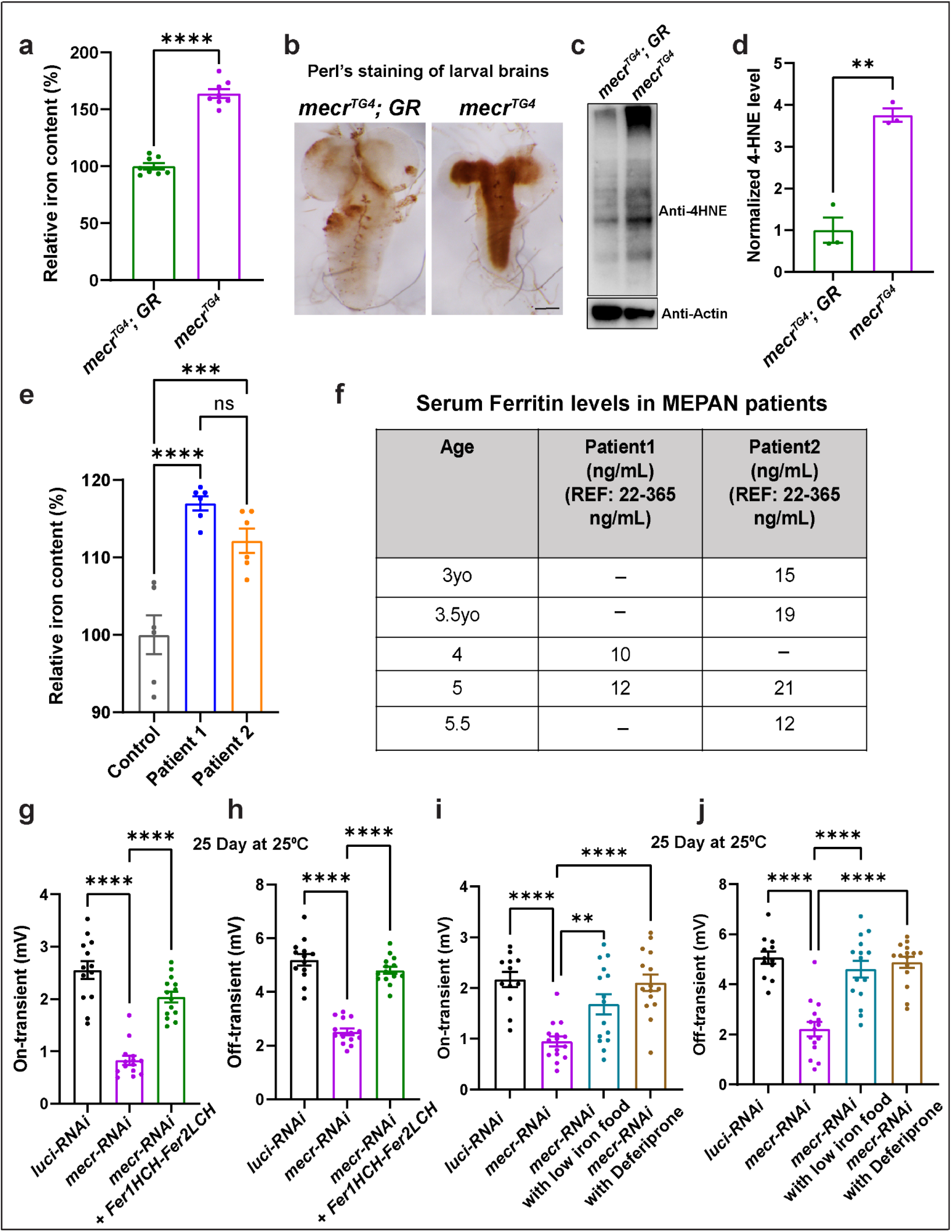
Loss of *mecr/MECR* impairs iron metabolism. (a) Relative amount of iron in whole body of *mecr^TG4^*mutants. (b) Iron staining in the larval brains of *mecr^TG4^* mutants. Scale bar: 2.54 mm. (c-d) Western blot and quantification showing 4-HNE levels in control and *mecr^TG4^* mutants. (e) Relative amount of iron in fibroblasts from patients and parental control. (f) Table showing patients’ clinical serum ferritin levels. (g-h) Quantification of ERG traces showing the effects of ferritin overexpression on synaptic transmission in 25-day-old flies with neuronal knockdown (*elav-GAL4*) of *mecr*. n = 13 (*luci-RNAi*), n = 14 (*mecr-RNAi*), n = 14 (*mecr-RNAi* + *Fer1HCH-Fer2LCH*). (i-j) ERG quantification showing the effects of low iron food and Deferiprone treatment on 25-day-old flies with neuronal knockdown (*elav-GAL4*) of *mecr*. n = 12 (*luci-RNAi*), n = 15 (*mecr-RNAi*), n = 15 (*mecr-RNAi* with low iron treatment), n = 15 (*mecr-RNAi* with low Deferiprone treatment). For statistical analyses between two samples, we performed two-tailed Student’s t-test, and for three samples, one-way ANOVA followed by a Tukey’s post-hoc test are performed. Error bars represent SEM (**p < 0.01; ***p < 0.001; ****p < 0.0001).

Iron is present in two interchangeable forms: ferric (Fe^3+^) and ferrous (Fe^2+^) forms. The conversion of (Fe^3+^) to (Fe^2+^) or (Fe^2+^) to (Fe^3+^) creates reactive oxygen species (ROS) via the Fenton reaction ^58, 59^. Increased ROS triggers lipid peroxidation and the generation of toxic hydroxynonenal (HNE)-modified protein adducts ^60–62^. Hence, we measured the levels of ROS using an antibody against 4-HNE, a marker of lipid peroxidation and oxidative damage ^58, 62, 63^. Indeed, we noted a three-fold elevation of the 4-HNE level in *mecr^TG4^* mutants (Fig. 6c- d). These data indicate that iron toxicity may lead to lipid peroxidation when *mecr* is lost. Alternatively, the impaired activity of Complex-I or Complex-III documented in Extended Data Fig. 6a may lead to increased ROS production.

Next, we evaluated iron levels in patient-derived fibroblasts. Compared to the control cells, fibroblasts from both patients displayed an elevated level of iron (Fig. 6e). Levels of ferritins, iron-binding proteins, in serum serves as another indicator of iron homeostasis in humans. Generally, low levels of serum ferritins indicate impaired iron metabolism and low iron storage ^64^. Interestingly, low levels of serum ferritin were noticed in both patients at different ages (Fig. 6f) and the levels of Ferritins are also comparatively low in another cohort of MEPAN patients (Table 1). The data indicate that loss of *mecr*/*MECR* impairs iron metabolism.

**Table 1.**
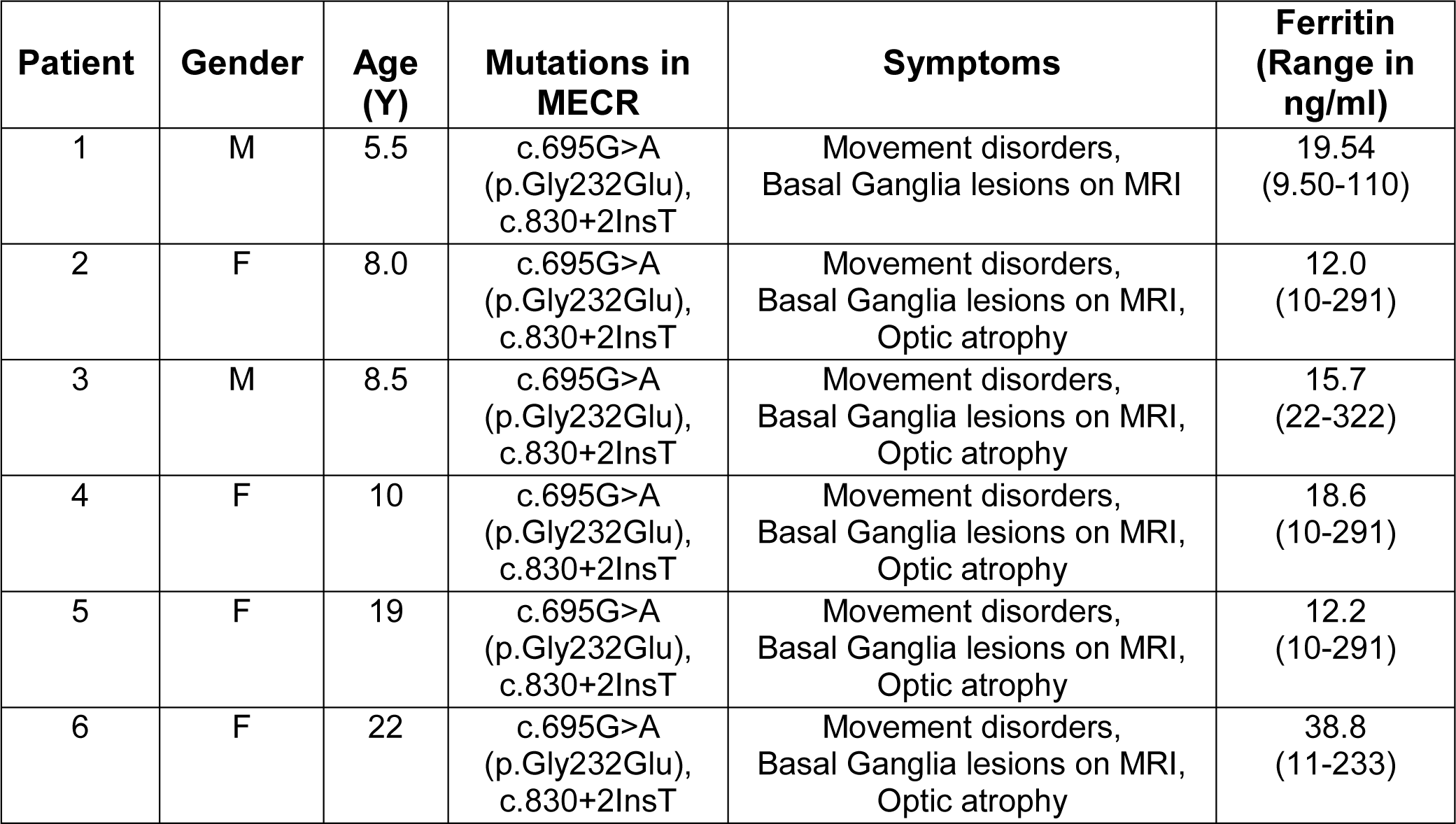
Detail of other MEPAN patients including Serum Ferritin levels. Table showing MEPAN patients including gender, age, mutations, symptoms, and ferritin levels in blood. Patient 3 was described by Heimer et al., 2016. Rest five are newly identified patients.

Since we noticed high iron levels in patient-derived fibroblasts and *mecr* mutants as well as low serum ferritin levels in MEPAN patients, we assessed if reducing iron levels can alleviate the neurodegenerative phenotypes in aged flies when *mecr* is reduced in neurons. First, we expressed Ferritin heavy chain (Fer1HCH) and Ferritin light chain (Fer2LCH), which form heteropolymeric complexes that chelate iron and reduce iron toxicity ^54, 65^. Increasing Ferritin levels rescued the ERG synaptic transmission defects as well as the climbing ability when *mecr* levels are reduced in neurons (Fig. 6g-h and Extended Data Fig. 7a-b). Second, we fed the flies an iron chelator, deferiprone ^66, 67^ or raised the flies on low iron food by replacing the iron-rich molasses with sucrose ^54^. Both treatments significantly improved the ERG and climbing defects (Fig. 6i-j and Extended Data Fig. 7c-d). These data argue that elevated iron play a role in the demise of neurons when *mecr* levels are reduced.

### Loss of Mecr affects Fe-S biogenesis

Impaired Fe-S cluster biogenesis in mitochondria have been reported to cause iron accumulation ^68, 69^. The Fe-S cluster biogenesis complex is composed of five proteins (Fig. 7a) including Iron-Sulfur Cluster Assembly Enzyme (ISCU), NFS1 Cysteine Desulfurase (NFS1), LYR Motif Containing 4 (LYRM4), Frataxin (FXN) and Acyl-ACP ^56^. Given the role of Acyl-(C14-C18)-ACP in Fe-S cluster biogenesis ^15, 70, 71^, we hypothesized that loss of *mecr* affects the length of the acyl chain and impairs iron-sulfur cluster biogenesis. We first assayed the activity of mitochondrial aconitase, which requires 4Fe-4S clusters for proper function and serves as a read-out for Fe-S cluster biogenesis defects ^72^. We find that the enzymatic activity of aconitase is significantly reduced in *mecr* mutants (>50%) (Fig. 7b) suggesting a defect in Fe-S cluster biogenesis. We then performed co-immunoprecipitation experiments to evaluate the integrity of the Fe-S cluster biogenesis complex upon loss of *mecr*. We immunoprecipitated either Nfs1 or Iscu and immunoblotted against Iscu or Nfs1 respectively using protein lysate from *mecr* null mutants. We observed that the interaction between Nfs1 and Iscu is impaired in the null mutants (Fig. 7c). No obvious difference in the levels of Nfs1 and a moderate reduction in the levels of Iscu protein was noted in the inputs (Fig. 7c). Hence, loss of *mecr* affects the assembly of Fe-S biogenesis protein complex and reduces Fe-S biosynthesis. We then assessed aconitase activity in patient-derived fibroblasts. Aconitase activity was also reduced in fibroblasts of patients compared to that of the control fibroblasts (Fig. 7h). Although the decrease in Aconitase activity is clearly less severe than in flies, this is not unanticipated as patient cells have residual MECR activity. Next, we performed co-immunoprecipitation to assess the interaction between ISCU and NFS1 using fibroblasts derived from the patients and parent control. However, we did not detect an impaired interaction between these two proteins (Extended Data Fig. 8), possibly because of residual MECR protein activity. Based on our observations, we propose that loss of *mecr*/*MECR* affects the Fe-S biogenesis.

**Figure 7:**
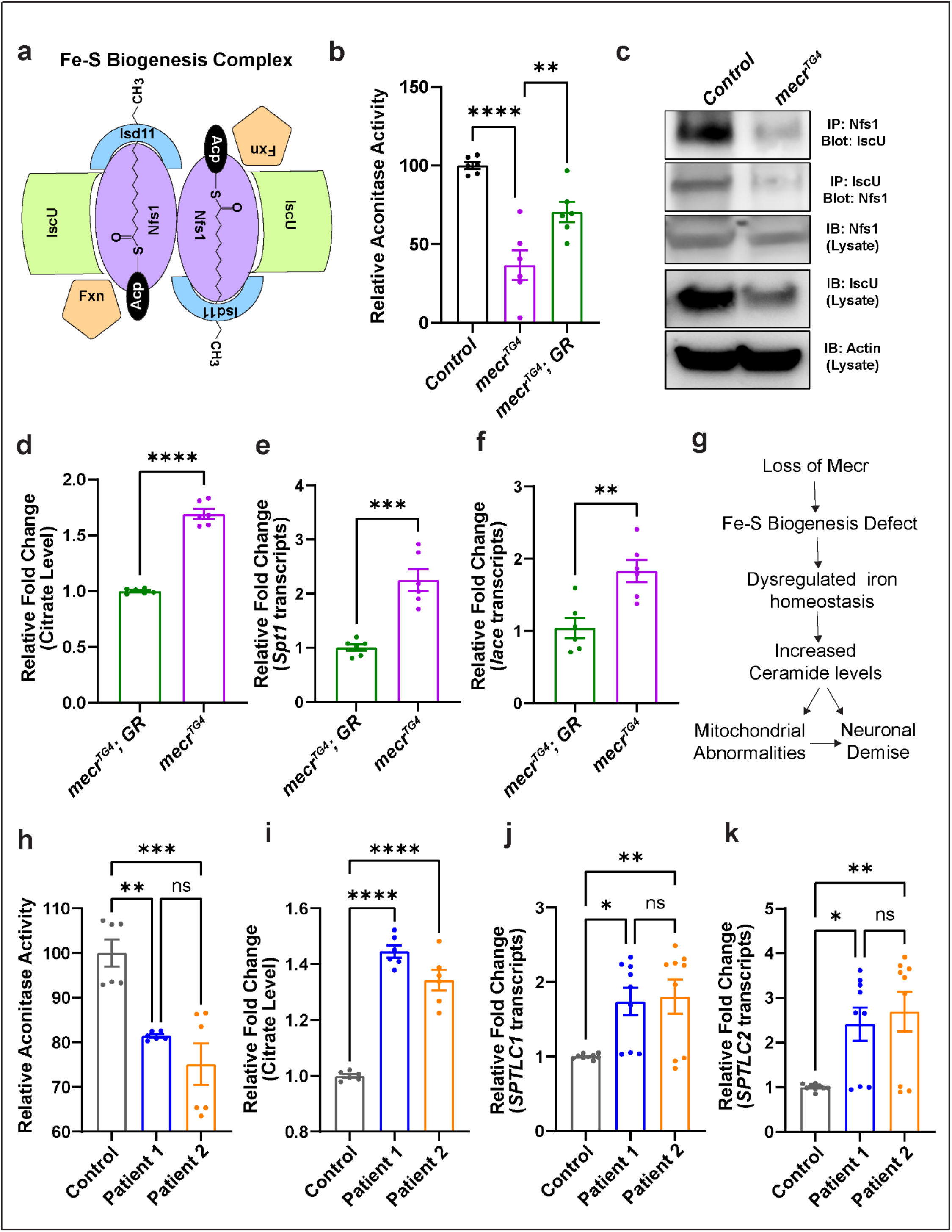
Loss of *mecr/MECR* reduces aconitase activity and promotes ceramide synthesis. (a) Diagram of the proteins of Fe-S biogenesis complex. (b) Relative aconitase activity in the *mecr^TG4^* mutants. (c) The interaction between Nfs1 and Iscu in the *mecr^TG4^* mutants is reduced. (d) Relative citrate levels in the *mecr^TG4^* mutants. (e-f) Relative fold changes in the transcript levels of *Spt1* and *lace* in *mecr^TG4^* mutants. (g) Proposed mechanism of neurodegeneration upon loss of *mecr*. (h) Relative aconitase activity in the fibroblasts from patients and parental control. (i) Relative citrate levels in patient-derived fibroblasts. (j-k) Relative fold changes in the transcript levels of *SPTLC1* and *SPTLC2* in patient-derived fibroblasts. For statistical analyses between two samples, two-tailed Student’s t test, and for three samples, one-way ANOVA followed by a Tukey’s post-hoc test are carried out. Error bars represent SEM (**p < 0.01; ***p < 0.001; ****p < 0.0001).

### Dyshomeostasis of iron metabolism leads to elevated ceramide levels

A reduction in aconitase activity has been reported to elevate citrate levels ^73–75^. We therefore assessed citrate levels and found that they are significantly increased in *mecr* mutants (Fig. 7d). Elevated citrate levels have previously been shown to cause an increase in palmitate and ceramides ^76, 77^. As we observe an upregulation in dihydroceramides (d18:0-16:0) (Fig. 3c and e), an increase in *de novo* synthesis of ceramides seems likely. We therefore assessed the transcript levels of the key rate limiting enzymes in the *de novo* synthesis of ceramides. As shown in Fig. 7e-f, there is a significant increase in transcript levels of *Spt1* and *lace* in *mecr* mutants. Hence, we argue that loss of *mecr* leads to dysregulated iron homeostasis, which in turn upregulates palmitate and ceramide levels (Fig. 7g). We also noticed increased citrate levels (Fig. 7i) as well as upregulated transcript levels of *SPTLC1* and *SPTLC2* in patient-derived fibroblasts (Fig. 7j-k). Overall, these data indicate that iron metabolism dyshomeostasis impairs aconitase activity to promote citrate levels and to upregulate ceramide synthesis.

## Discussion

Despite the discovery of mtFAS in the late 1980s ^78, 79^, its contribution to lipid homeostasis has not been systematically investigated *in vivo* in multicellular organisms. The recent identification of loss-of-function mutations in *MECR* as the cause of MEPAN syndrome, a pediatric-onset neurodegenerative disorder^10^, highlights the importance of mtFAS in neuronal maintenance in humans. To investigate if and how impaired mtFAS affects lipid homeostasis and leads to neurodegeneration, we generated a fly mutant and characterized MEPAN patient-derived fibroblasts. In flies, severe loss-of-function alleles of *mecr* cause lethality, whereas neuronal knock-down of *mecr* leads to age-dependent locomotor defects, visual impairments, and progressive photoreceptor degeneration. Our comparative analyses between flies and patient fibroblasts revealed elevated ceramide levels and mitochondrial dysfunction when *mecr/MECR* is lost or reduced. We propose that loss of *mecr*/*MECR* affects the Fe-S biogenesis, possibly because the ACP-acyl carbon chain length is severely reduced, which leads to elevated cellular iron level. Reducing the levels of either iron or ceramide strongly suppresses the age-dependent phenotypes in flies. These data argue that low levels of Fe-S clusters, elevated iron and/or increase ceramide play an active role in the demise of neurons when *mecr/MECR* levels are reduced. Moreover, they also point to the functional relationship between these two metabolic pathways implicated in neuronal maintenance.

To our knowledge, *mecr*/*MECR* has not been previously implicated in ceramide or iron metabolism. The *de novo* synthesis of ceramides occurs in the endoplasmic reticulum (ER) whereas acidic and neutral sphingomyelinases break down sphingomyelin to produce ceramides in lysosomes and at the plasma membrane respectively^80^. Our data suggest that the ceramides and iron metabolomic defects are connected and that *MECR/Mecr* plays a central role in maintaining homeostasis between these pathways by regulating mtFAS. Elevated ceramide levels impair membrane dynamics by reducing the fluidity of membranes and trafficking of membrane compartments in the cell ^44, 80–83^. They also result in loss of mitochondrial membrane potential, impaired ETC activity, and formation of ceramide channels on the outer mitochondrial membrane, eventually causing cell death ^84, 85^.

Membrane and mitochondrial defects have been shown to induce the demise of neurons in neurodegenerative disorders such as Parkinson’s disease (PD) and Friedreich ataxia (FRDA) ^53, 54, 86, 87^. A partial loss of *FXN* causes FRDA ^67, 88^. The Frataxin protein is a key component of the Fe-S cluster biosynthesis complex which also contains acyl-ACP. Loss of *mecr* (this work) or loss of *fh* ^89^ affects the Fe-S cluster synthesis as evidenced by reduced activity of aconitase. We previously showed that loss of Frataxin in flies as well as in cardiomyocytes of FRDA patient causes increased levels of dhSph, Sph, dhCer, Cer ^53, 54^ and increased ceramide levels have also recently been documented in FRDA patient-derived fibroblasts ^88^. The loss of Fe-S cluster synthesis should lead to an elevation of iron, which is also observed upon loss of *fh* ^53, 54^ and *mecr* (this work). Hence, elevated iron may play a role in the pathogenesis. Indeed, lowering the iron levels in both models is beneficial, suggesting that the Fe-S cluster defects may not account for all the observed phenotypes unless both insults, low Fe-S cluster synthesis and high iron synergize to cause severe cellular defects. Restoring one insult may therefore be sufficient to partially suppress the observed phenotypes.

A Fe-S cluster synthesis defect, as well as a reduction in aconitase activity and elevated iron levels has also been documented in flies that lack *pink1* ^90^. Loss of PINK1 causes parkinsonism in human ^91^ and fibroblasts derived from patients as well as *pink1* mutant flies exhibit an accumulation of ceramides ^42^. Elevated levels of iron and ceramides have also been associated with PD ^86, 87, 92–94^ and iron deposition has been reported in the substantia nigra of the postmortem PD brain ^95–97^. Finally, elevated levels of ceramide in plasma, serum, and postmortem CSF have been reported in PD patients ^92^. Based on these data, we argue that the pathway studied here is relevant to at least some forms of PD or parkinsonism.

The elevated levels of iron may lead to the upregulation of ceramides through different mechanisms. We noted an elevation in citrate levels as well as an upregulation in the transcript levels for the proteins required for the *de novo* synthesis of ceramide. Previously another study reported citrate accumulation in HeLa cells upon knockdown of ACP ^23^. A significant increase in citrate due to reductive carboxylation was also observed in mtFAS deficient C2C12 cells ^6^. Elevated citrate levels inhibit fatty acid transport to mitochondria for beta-oxidation and increases palmitoyl-CoA availability for *de novo* ceramide synthesis ^76, 77^. In addition, other mechanisms facilitating ceramide accumulation may also be operative due to iron metabolism impairments. For example, impaired iron metabolism produces ROS ^58, 59^, which can enhance the activity of both neutral and acidic sphingomyelinases ^98, 99^. Alternatively, the impairment in Fe-S cluster biogenesis may also lead to elevated ceramide levels. Fe-S clusters are required for the activity of Lipoic Acid Synthase (LIAS) to produce Lipoic acid, which is known to inhibit neutral SMases ^100–102^. Moreover, mtFAS produces octanoate, the substrate for LIAS. Hence, reduced lipoic acid production may also lead to the activation of neutral SMases. However, given that we observe an elevation of dihydroceramides and *de novo* synthesis, the latter two hypothesis seem less likely or may only play a minor role.

It is worth noting that although the effects of *mecr* loss-of-function in flies are mostly consistent with the phenotypes we observe in patient-derived fibroblasts, the fly phenotypes are typically more pronounced. We argue that the main reason for this difference is the partial loss of *MECR* function in patients whereas flies carry null alleles. Indeed, expression of some *MECR* variants from MEPAN patients in a fly mutant background suggests that the variants that affect single amino acids are relatively weak hypomorphic alleles whereas the truncating variant is a strong loss-of-function allele. Indeed, 7 out of 8 patients ^10, 11^ carry at least one allele that is a weak hypomorph (Extended Data Fig. 2b). We argue that stronger allelic combinations in humans may result in embryonic lethality similar to complete loss of Mecr function results in embryonic lethality in mice ^4^ and larval lethality in flies (this study). Hence, the fly phenotypes are likely to be more severe than the phenotypes in patient fibroblasts.

A role for mtFAS in Fe-S cluster biogenesis has been recently reported in yeast as well as in vertebrate cell culture studies ^15, 70, 71^. ACP with an acyl chain length of 14 to 18 carbons (Acyl-ACP) interacts with ISD11 and facilitates the assembly of the Fe-S cluster biogenesis complex ^15, 71^. However, whether a reduction in acyl chain length affects Fe-S cluster biogenesis was not tested. Our data show that the five-protein complex is not properly assembled in *mecr* null mutants. However, in patient fibroblasts, we did not detect an obvious defect in the complex assembly, possibly because some Acyl-ACP is still produced. However, other data are consistent with impaired Fe-S biogenesis. A defect in Fe-S biogenesis affects the activity of aconitase, a mitochondrial enzyme, which requires [4Fe-4S] clusters ^72^. A previous study in mice used a histochemical approach to measure aconitase in Mecr conditional KO Purkinje cells and did not detect a difference in aconitase activity ^9^. Yet, our data show that there is a Fe-S biogenesis defect in the fly *mecr* mutants as well as the patient-derived fibroblasts based on aconitase activity. Moreover, our data show that *mecr* mutants and patient-derived fibroblasts have elevated iron levels. Dysregulated iron metabolism has not yet been reported in patients with MEPAN syndrome. However, in both patients reported in this paper, we noticed low plasma Ferritin levels, as well as high cellular iron levels and impaired aconitase activity in fibroblasts. We recently identified five more MEPAN patients (Table1). All patients exhibit low plasma levels of Ferritin, but most are still within a borderline normal range (Table1). Interestingly, mRNAs encoding Ferritins contain an Iron Responsive Element (IRE) in the 5’ UTR of the mRNA. In the presence of low Fe-S clusters, the IRP1 protein binds to the 5’UTR IRE of the *ferritin* mRNA and reduces its translation leading to reduced plasma Ferritin levels ^64, 103^. Hence, the low plasma Ferritin levels in patients provides further evidence for a defect in iron metabolism. Reduced Ferritin levels may also cause increased iron deposition in the cells leading to neurodegeneration as seen in Neuroferritinopathy ^104–106^. Interestingly, Patient 3 (Table1), who exhibits the most severe phenotype including manifestation of symptoms at the youngest age and quickly deteriorating symptoms, has significantly elevated iron levels (159 µg/dL, Range: 50 to 120 µg/dL) and Ferritin levels below the normal range (Table1). Finally, the data are also consistent with the observation that increasing Ferritin levels in flies alleviate neurodegenerative phenotypes associated with *mecr* knock-down in adult flies. Hence, we propose that impaired Fe-S biogenesis results in increased iron and ceramide levels leading to age-dependent impairment of neuronal function in *mecr* mutant flies and MEPAN patients.

## Acknowledgments

We extend our thanks to the affected individuals and families who participated in this study. We thank Teresa M. Dunn, Danny Miller, Susan J. Hayflick, Alexander J. Kastaniotis, Maimuna S. Paul, and Hyunglok Chung for helpful discussion, Hongling Pan and Wen-Wen Lin for injections to create transgenic flies, and Jinyong Kim for Mass spectrometric analyses. Our sincere thanks to Fanis Missirlis (*UAS-Fer1HCH*, *UAS-Fer2LCH* stock), the Bloomington Drosophila Stock Center, the Vienna Drosophila Resource Center, and the Kyoto Drosophila Genetic Resource Center and the Developmental Studies Hybridoma Bank from the University of Iowa for providing fly stocks and reagents. We acknowledge the Shan and Lee-Jun Wong fellowship from Baylor College of Medicine to D.D. We thank the BCM Intellectual and Developmental Disabilities Research Center (IDDRC) confocal microscopy core, supported by the National Institute of Child Health & Human Development (U54 HD083092). H.J.B. is supported by the NIH Common Fund, through the Office of Strategic Coordination/Office of the NIH Director (U54NS093793), NIH/ORIP (24OD022005 and R24OD031447) and is a recipient of the Chair of the Neurological Research Institute. The content is solely the responsibility of the authors and does not necessarily represent the official views of the National Institutes of Health.

## Author Contributions

Conceptualization, D.D. and H.J.B.; Investigation, D.D., O.K., P.C.M., S.K.B., Z.Z., J.H.P, R.V.S., G.L., J.N.K., M.T.W., G.H., and M.G; Resources, H.J.B., A.P., B.A.K.; Writing – Original Draft, D.D., H.J.B.; Writing – Review & Editing, D.D., O.K., P.C.M., G.L., S.K.B., J.N.K., M.T.W., J.H.P, R.V.S., A.P., B.A.K., G.H., and H.J.B.; Funding Acquisition, H.J.B; Supervision H.J.B.

## Ethics statement

The fibroblasts were obtained with patient consent approved by the Institutional Review Board.

## Data and reagent availability

All the data and reagents used in this study are available upon reasonable request to the corresponding author.

## Materials and Methods

### Fly Stocks maintenance and survival assay

Flies were maintained in a standard fly food at room temperature and the experiments were carried out in 25°C unless otherwise noted. The stocks used in this work were either generated either in the lab or obtained from other labs and stock centers including Bloomington Drosophila Stock Center (BDSC) or Vienna Drosophila Resource Center (VDRC). *yw* was used as the ‘control’ for most of the experiments. For the lifespan assay, newly emerged flies were collected in fresh vials, and approximately, 10-15 flies were kept in each vial from each genotype at 25°C with 12h dark/12h light conditions. The flies were transferred into a new vial every 2-3 days followed by documenting the number of dead flies. The significance of survivability was calculated using Log-rank (Mantel-Cox) test and Gehan-Beslow-Wilcoxon test in GraphPad Prism software.

### Generation of CRISPR mutant

We generated *mecr^TG4^* mutant flies as described previously ^24^. In brief, we designed sgRNAs targeting the coding intron of CG16935/*mecr* (TATGCAGTTCATGCCGGTTATGG) as well as the 200 bps left and right homology arms. A construct containing the homology arms and sgRNA sites to linearize the homology donor construct is synthesized in pUC57_Kan_gw_OK vector by Genewiz (South Plainfield/NJ). sgRNA targeting mecr is cloned in pCFD3 vector ^107^. The gRNAs, homology arms, and the cassette containing *attP-FRT-SpliceAcceptor-T2A- GAL4-polyA-3XP3GFP-polyA-FRT-attP* sequence was subcloned in the homology intermediate vector from pM37 vector, creating the homology donor vector ^25^. sgRNA vector and homology donor vector are injected in nosCas9 expressing embryos as described in Kanca et al. 2019^108^. Eclosed flies are crossed with *yw* flies, and the 3XP3-GFP positive progeny were selected on the basis of GFP expression in the adult eyes.

### Cloning and transgenics

The *MECR* (NM_016011.2) human cDNA was cloned into the destination vector, pGW-attB-HA as previously described ^109^. Briefly, cDNA in the pDONR221 vector was shuttled to the pGW-attB-HA using Gateway cloning (LR clonase II, Thermo Fisher Scientific). For generating the variants, the Q5 site-directed mutagenesis (NEB) in the pDONR221 vector was performed using the following primers MECR^Gly232Glu^, For 5’- AAGAGTCTGGAGGCTGAGCAT-3’, Rev 5’-CAGTCTGTCACTCAGCTTC-3’; MECR^Arg258Trp^, For 5’- GCCCCAGCCATGGCTTGCTCT-3’; Rev 5’-ATGTCCTTAAAGAAGTTTTTCATTTCGGGCC-3’; MECR^Trp285*^ For 5’-TGGTAACCTAGGGGGGGATGG-3’, Rev 5’-TGGTTCCTCCACGCGCTA-3’. All variant sequences were Sanger verified after LR reaction to the destination vector.

For, the fly *UAS-mecr-HA* line, the clone from DGRC (Cat – 1660183) was obtained. As previously described ^110^, flanking Gateway compatible attB sites were inserted by PCR to the *mecr* template and shuttled to the pDONR223 by BP clonase II (Thermo Fisher Scientific) using the following PAGE purified primers: mecr_attB_For 5’- GGGGACAAGTTTGTACAAAAAAGCAGGCTTCACCATGTTGCGCAGAGGCTTTTTATCTC-3’ and mecr_attB_Rev 5’- GGGGACCACTTTGTACAAGAAAGCTGGGTCCTAAATGCTCATATCCAGTATGTAC-3’. *yw ΦC31 integrase; VK33 (PBac{y[+]-attP}VK00033)* flies were used for injecting the UAS constructs. For the *mecr-GFP* line, we obtained the FlyFos construct (CBGtg9060C0290D) from the SourceBiosciences and prepared the DNA for injection into the y1 w*, P{nos-phiC31\int.NLS}X; PBac{y+-attP-3B}VK00033 flies.

### Quantitative Real-Time PCR

Real time PCR was performed as described previously ^111^. In short, RNA was isolated from 2^nd^ larva of the indicated genotypes using the RNeasy Plus Mini Kit (Qiagen) followed by preparation of cDNA using 5× All-In-One RT MasterMix (Applied Biological Materials Inc.) as per the manufacturer’s protocol. Master Mix for the quantitative polymerase chain reaction was prepared using 2X Fast SYBR Green Master Mix (Thermo). Following primers were used for *mecr*: 5’-ATTCTGGCAGCTCCCATTAAC-3’ (Forward, primer set 1), 5’- CGGCTGGAAACTTGGGCTT-3’ (Reverse, primer set 1), 5’-CGTGGGCGACAAAGTCAAAG-3’ (Forward, primer set 2), 5’-CAACCTTCTTGGACACGATCA-3’ (Reverse, primer set 2); *Spt1*: 5’- ACCCTACTGCTCATAACCGTG-3’(Forward primer), 5’-GCGATTATTCGGTCTTCCTCCTC-3’ (Reverse primer); *lace*: 5’-CCGCGTACACTGAAATTCGC-3’ (Forward primer), 5’-CCGGATGGTAGTTGATCGAGC- 3’ (Reverse primer); *SPTLC1*: 5’-GGTGGAGATGGTACAGGCG-3’ (Forward primer), 5’- TGGTTGCCACTCTTCAATCAG-3’ (Reverse primer); *SPTLC2*: 5’-TGGGTTCCTACAACTATCTTGGA-3’ (Forward primer), 5’-CATACGCCATAGCAGCTTCTAC-3’ (Reverse primer). *rps13* was used as internal control for the fly genes and *18s rRNA* was used as internal control for the human genes. Reactions for each genotype was set up in triplicate using Applied Biosystems QuantStudio 6 machine. Relative fold change of the *mecr* transcript was calculated using 2^−ΔΔCq^ method and the graph was prepared using GraphPad Prism software.

### Fibroblast culture

Fibroblasts derived from the parent (control) and both patients with MEPAN syndrome were obtained from Undiagnosed Diseases Network and cultured in Dulbecco’s Modified Eagle Medium (Gibco, USA) supplemented with 10% fetal bovine serum (Invitrogen, USA) and 1% Antibiotic-Antimycotic (Gibco, USA). All cells were grown in 5% CO2 at 37°C. Harvested cells were washed with phosphate-buffered saline and stored as pellets at - 80°C until further analysis. All experiments were performed using the same passage number.

### S2 cell culture and transfection and immunostaining

S2 cells were cultured on polyD-Lysine coated coverslips in 24 well plates at room temperature (23°C) using Schneider’s Drosophila medium (Life Technologies) supplemented with 10% fetal bovine serum (FBS) and Penicillin-Streptomycin (Sigma). Transfection of S2 cells were carried out with 0.2ug DNA using Qiagen Effectene Transfection Reagent as per the manufacturer’s recommendation. After 48 hours of transfection, the cells were used for immunostaining. Briefly, the media was replaced with 500 μL of 4% PFA (paraformaldehyde) for 30 min at room temperature. Then, the coverslips were rinsed with 1X PBS for three to four times followed by treatment with 200 μL of 0.2% PBT for 15 min and blocking with 1% NGS for 1 hour. Subsequently, primary antibody incubation was performed at 4°C overnight. The next day, cells were washed with PBS 4 times, 5 min each, and incubated with respective secondary antibodies and DAPI for 60-90 min at room temperature. Finally, the coverslip with cells was briefly washed with 1X PBS and mounted using vectashield (Vector Labs, Burlingame, CA). The slides were either imaged immediately using a Leica sp8 (Leica) confocal microscope. The images were processed using ImageJ and Adobe Photoshop.

### Immunostaining of fly tissues

Immunostaining was performed as described previously ^109, 111^. Larvae were dissected in cold PBS followed by fixation in 4% PFA for either 20 min (salivary gland and fat body) or overnight (brain). Next, the tissues were washed briefly in PBST 3-4 times for 10-15 min each, blocked in 5% NGS and kept in primary antibody for overnight at 4°C. The following dilutions of primary antibodies were used: Rat Anti-Elav (1:500), Mouse Anti-Repo (1:50), Rabbit Anti-MECR (1:100), Mouse Anti-ATP5α (1:500), Mouse Anti-Cer (1:15). The next day, after washing with PBST 3-4 times for 15 min each, the tissues were incubated in secondary antibody in 5% NGS and PBT for 90 min at room temperature. All the secondary antibodies were used at a 1:200 dilution. The secondary antibody was removed followed by three washes with PBST 3-4 times for 15 min each. Finally, the tissues were mounted in vectashield (Vector Labs, Burlingame, CA) or RapiClear mounting medium. For the adult brains, 2% PBT was used for all the washes and the antibody incubations were performed for 48 hours. Imaging was performed by using either LSM880 (Zeiss) or Leica sp8 (Leica) confocal microscopes. The images were processed using Zen Blue (Zeiss LSM Image Browser), ImageJ and Adobe Photoshop.

### Electroretinogram

Electroretinograms were performed as described by ^109^. The flies were glued to the glass side and kept in complete darkness for at least one minute followed by inserting two glass electrodes: the ground electrode was placed onto the thorax and the recording electrode was placed onto the eye. The eyes were exposed to a pulse of light for 1 sec. The electroretinogram traces were recorded and data analyses were performed using LabChart8 Reader software.

### Climbing assay

Climbing assay was performed as described previously by ^111^. Briefly, 3-5 flies were taken in a plastic food vial and tapped 2-3 times onto the bottom. Next, two parameters were measured: first, the percentage of flies, who were able to climb ∼8 cm in 30 sec. Second, the average time taken by the flies to reach ∼8 cm distance within 30 sec.

### Transmission Electron Microscopy

Transmission electron microscopy was performed using retina from the adult fly eye as well as fibroblasts from patients and parental control. The samples were fixed using 4% PFA, 2% glutaraldehyde, and 0.1 M sodium cacodylate (pH 7.2) at 4°C for 48 hours followed by a secondary fixation in 1% OsO4. Subsequently, the samples were dehydrated by treating them with ethanol grades and propylene oxide. For this, a Ted Pella Bio-Wave microwave oven with vacuum attachment was used. Next, the samples were embedded in Embed-812 resin (Electron Microscopy Science, PA) for 72 hours. Then, ultra-thin sectioning was performed using a Leica UC7 microtome followed by staining with 1% uranyl acetate and 2.5% lead citrate. Finally, imaging was performed using a JEOL JEM 1010 transmission electron microscope with an AMT XR-16 mid-mount 16 megapixel camera.

### Lipidomic analyses

Lipidomic analyses was performed using the *mecr^TG4^* mutant larva and human fibroblasts.

#### Lipid extraction

Lipids were extracted from wild-type, mutant and rescue larva, as well as control and patient fibroblast pellets using a neutral and acidic lipid extraction as described previously ^112–114^. Briefly, deuterated internal standard mixture, described in the previous method ^114^ was added with chloroform:methanol (1:2, v/v) to all samples equally. After the addition of internal standards, the samples were tip-sonicated to disrupt the cells and extraction solvent was added to extract lipids while removing proteins and polar metabolites. Extracted lipids were stored at -80°C until mass spectrometry analysis.

#### Liquid chromatography-tandem mass spectrometry for lipidomics

The larva and human fibroblast lipid extracts were analyzed on a Hypersil GOLD Vanquish C18 UHPLC column (150 × 2.1 mm, C18 1.9 μm and 175 Å) using a Fourier transform Orbitrap Fusion Tribrid ID-X mass spectrometer (Thermo Fisher Scientific, USA) coupled to Vanquish Horizon UHPLC (Thermo Fisher Scientific, USA). Untargeted lipidomics was carried out as previously described, with modifications ^113, 114^. Briefly, a binary gradient at a flow rate of 300 µL/min using aqueous phase (water:acetonitrile=6:4, v/v) and organic phase (isopropanol:methanol:acetonitrile=8:1:1, v/v/v) with 10 mM ammonium formate and 0.1% formic acid as modifiers was applied to separate the lipids. Organic phase was increased from 30% to 95% over 15 min, maintained at 95% for 3 min and equilibrated to 30% for 5 min for the next injection. The analytical column was maintained at 50°C. A full scan MS (250–1600 m/z in positive ion mode and 350-1700 m/z in negative ion mode) was acquired in the Orbitrap with a resolution of 60,000 at m/z 200. Injection time was set at 50 ms in MS and dynamic exclusion was enabled with an 8 s exclusion duration time. MS/MS spectra were obtained at a resolution of 15,000 at m/z 200 with injection time at 60 ms. MS/MS scans were acquired for 1.5 sec, followed by a MS scan. In positive ion mode, spray voltage of 3.5 kV and stepped collision energy of 30, 35 and 40% in higher-energy collisional dissociation (HCD) were applied. Collision-induced dissociation was triggered only for ions with detection of m/z 184 in HCD fragmentation. In negative mode, spray voltage of 3 kV and stepped collision energy of 28, 32 and 40% in HCD were used.

#### Mass Spectrometry data analysis for lipidomics

The acquired tandem-mass spectra were processed using LipidSearch 4.2 (Thermo Fisher Scientific, USA) to annotate lipids. Lipids were annotated by comparing precursor ion masses and corresponding MS/MS spectra against the database. Precursor ion mass tolerance was set to 5 ppm and fragment ion tolerance was set to 8 ppm. Annotated lipids were quantified by calculating peak areas using PyQuant, based on the precursor ion masses and retention times ^115^. All lipids were normalized by peak area of internal standard sharing the identical head group of lipids.

### ATP determination

ATP determination was performed using ATP determination Kit from Thermo (A22066) as per the manufacturer’s instruction. 25 larvae were homogenized in 100 µL of homogenization buffer followed by boiling for 5 min and centrifugation at 15000 rpm for 3 min at 4°C. The supernatant was collected in a separate tube. For the fibroblasts, ∼1×10^4^ cells were washed twice with PBS followed by centrifugation to make a cell pellet. The pellet was resuspended with ATP releasing agent from Sigma (FLSAR-1VL). 10 µL of the supernatant was diluted with 90 µL of homogenization buffer and used for further experiments. Finally, the diluted sample was added into Luciferase-luciferin mix (1:10) followed by measurement of luminescence. Total protein levels were determined and used to finally normalize the data.

### Mitochondrial Membrane potential analyses

Mitochondrial membrane potential was performed as described previously ^37^. Briefly, larvae were dissected in DSM and stained with 100 nM TMRE for 20-30 min followed by two brief washes in PBS. Samples were mounted in 1X PBS and immediately imaged using 63X objective of Zeiss880. The images were processed using Zen Blue (Zeiss LSM Image Browser) and Adobe Photoshop software.

### ETC activity

Activities of ETC complexes were measured by kinetic spectrophotometric assays as previously described ^116^. Briefly, 150 second instar larva were homogenized in 500 µL of Mitochondria Isolation Buffer (225mM Manitol, 75mM Sucrose, 10mM MOPS, 1mM EDTA, 2.5mg/ml BSA in milli-Q water). After homogenization, protein concentration was determined by Bradford assay. Enzymatic activity of the individual OXPHOS complexes was measured using a Tecan Infinite M200 microplate plate reader. The assay is based on the measurement of oxidation/reduction of substrates or substrate analogues of individual complexes. The enzyme activity of individual complexes was normalized to the protein concentration.

### Seahorse assay

The oxygen consumption rates of fibroblasts were measured using the XF24 extracellular flux analyzer from Seahorse Bioscience, Agilent as described previously ^50^. Briefly, the cells were plated in 4 replicates with ∼60000 cells in each well and the XF24 cartridge was equilibrated using calibration solution at 37°C for overnight. The next day, the media was replaced with XF assay medium (5mM glucose/5mM glactose, 2mM pyruvate in DMEM media, pH 7.0) and different mitochondrial stressors, namely Oligomycin (500 nM), FCCP (500 nM), Rotenone and Antimycin A (100 nM each) were added to the wells containing the fibroblasts in a sequential manner. A protocol of 3 min mixing followed by 3 min measurement and 2 min incubation time was followed to measure the oxygen consumption rates. Different respiratory parameters were calculated by using Seahorse Wave software. Finally, the protein concentrations of each well were measured and used to normalize the readings.

### Ferrozine assay

Ferrozine assay was performed following the slightly modified protocol from Missirlis et al., 2006; Schober et al., 2021^117, 118^. Briefly, 10 larvae or cell pellets were homogenized in lysis buffer followed by centrifugation at 16000xg for 10 min. Concentrated HCL was added to the supernatant followed by heating for 20 min at 95°C. Then, the tube was centrifuged at 16000xg for 2 min. 20 µL of 75 mM Ascorbate and 20 µL of 10 mM Ferrozine were added to 50 µL of supernatant and mixed. Finally, 40 µL of saturated ammonium acetate was added to the mixture followed by absorbance measurement at 560nm using FLUOstar OPTIMA microplate reader (BMG Labtech). Results were normalized with the protein concentrations.

### Iron staining

Iron staining was performed as described previously ^54^. In short, larval brains were dissected in ice-cold PBS followed by fixation in 4% PFA for 20 min. Subsequently, the brains were washed quickly in 0.4% PBT and incubated in Pearl’s solution (1% K4Fe(CN)6 and 1% HCL in PBT) for 10-15 min. After incubating with Pearl’s solution, brains were again washed quickly with 0.4% PBD and incubated for 3-5 min with DAB solution (10 mg DAB in 0.07% H2O2 and 0.4% PBT) for enhancing the signal. Lastly, the brains were washed briefly in 0.4% PBT, mounted in (Vector Labs, Burlingame, CA) and imaged with a Zeiss SteREO Discovery.V20 microscope equipped with Omax 18.0 MP Camera. Images were processed using ImageJ and Adobe Photoshop software.

### Aconitase assay

Aconitase assays were performed using Aconitase Activity Assay Kit from Sigma (MAK051) ^119^. Briefly, ∼50 larva or fibroblast pellets were homogenized in 100 µL ice-cold assay buffer and the homogenate were centrifuged at 800xg for 10 min. 65 µL of supernatant taken in a fresh tube and centrifuged at 20000xg for 15 min. Pellets were resuspended in 100 µL assay buffer and used for the subsequent steps following manufacturer’s protocol. The absorbance was measured using FLUOstar OPTIMA microplate reader (BMG Labtech). Finally, the readings were normalized with protein concentrations.

### Citrate Assay

Citrate assays were performed using Citrate Assay Kit from Sigma (MAK057). In brief, ∼50 larva or fibroblast pellets were homogenized in 100 µL assay buffer followed by centrifugation at 15,000xg for 10 minutes at 4°C. Supernatant was used for the subsequent steps following manufacturer’s protocol with slight modifications. After 15 min incubation, absorbance was measured using FLUOstar OPTIMA microplate reader (BMG Labtech). Finally, the readings were normalized with protein concentrations.

### Protein Isolation and Western Blot

Larvae or cell pellets were homogenized in ice-cold RIPA buffer with 1X liquid protease inhibitor (Gen DEPOT) followed by centrifugation for 10 min at 13000 rpm at 4°C. The supernatant was collected in a separate tube, where 4X Laemmli Buffer containing β-mercaptoethanol was added. After mixing well, the samples were heated at 95°C for 10 min followed by keeping them in ice until loaded. The proteins were separated using 4–20% gradient polyacrylamide gels (Bio-Rad Mini-PROTEAN® TGX™) at 90-100 volts. After electrophoresis, western blot was performed using a polyvinylidene difluoride membrane (Immobilon, Sigma) followed by blocking in either skimmed milk or 5% BSA for 60-90 mins. Finally, the membranes were incubated in the primary antibody overnight. The following dilutions were used for the primary antibodies: Anti-Iscu (1:1000), Anti-NFS1 (1:1000), Anti-Actin (1:10,000). The next day, the membrane was washed briefly followed by incubation in horseradish peroxidase-conjugated secondary antibody in blocking solution (1:5000) for 90 min at room temperature. Next, the membrane was washed three times and Fixer-Developer (1:1) solution from Western Lightning Chemiluminescence Reagent Plus (PerkinElmer, NEL104001EA) enhanced chemiluminescence (ECL) was used to develop the signal, which was visualized using Bio-Rad ChemiDoc MP Imaging System.

### Co-Immunoprecipitation

Co-immunoprecipitation was performed as described previously ^120^. Briefly, the protein samples from larvae or fibroblasts containing 1-2 mg proteins were mixed with Protein A/G beads (SCBT) and the respective antibodies in an end-over-end rotator at 4°C overnight. The next day, the samples were briefly centrifuged and washed in IP buffer three to four times followed by denaturation, PAGE separation and western blotting.

### Preparation of low iron fly food and drug administration

For preparing the low iron food for flies, dry yeast (10% w/v), agar (0.6% w/v) and sugar (10% w/v) were mixed in mili-Q water, microwaved and aliquoted in vials. After the food was solidified, the vials were used immediately or stored at 4°C for future use. For testing the effects of reduced ceramide or iron levels, fly food with different drugs was prepared and administered along with corresponding control food. Myriocin (100 mM in ethanol), Desipramine (1.126 mM in milli-Q water) and Deferiprone (163 uM in milli-Q water) were mixed in regular food and used immediately or stored at 4°C for future use. For all the experiments, the flies were transferred into a fresh vial in 2 to 3-day intervals.

### Statistical analyses and data quantification

The data used for quantification were obtained in at least three biological replicates, and GraphPad Prism (GraphPad Software Inc., CA, U.S.) was used for the statistical analyses and quantification. For comparing two groups, unpaired t-test (two-tailed) and for comparing more than two groups, analysis of variance (ANOVA) was used. All the data are represented as bar plots and the error bar represents ± SEM. P-value less than 0.05 is considered significant (*P < 0.05, **P < 0.01, ***P < 0.001, and ****P < 0.0001) and less than 0.05 is considered nonsignificant (ns).

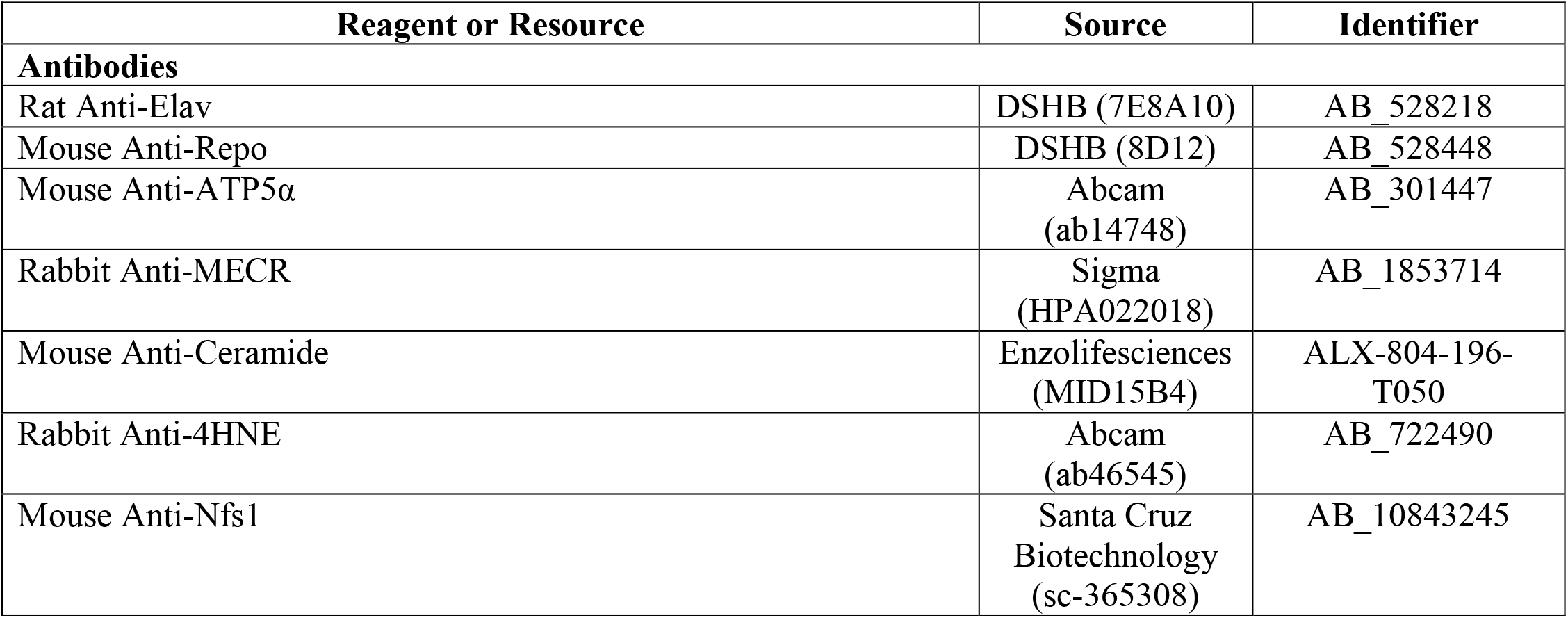
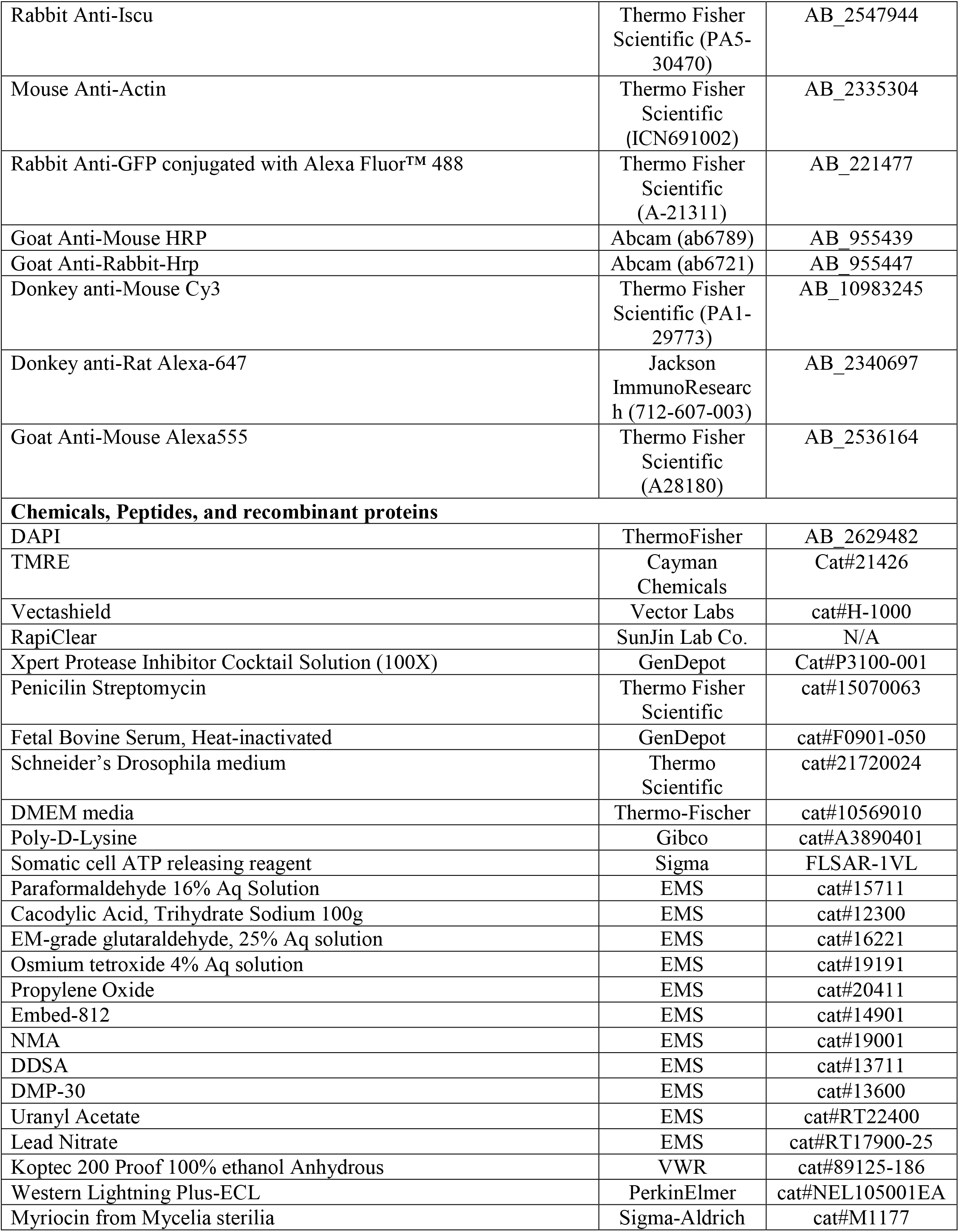
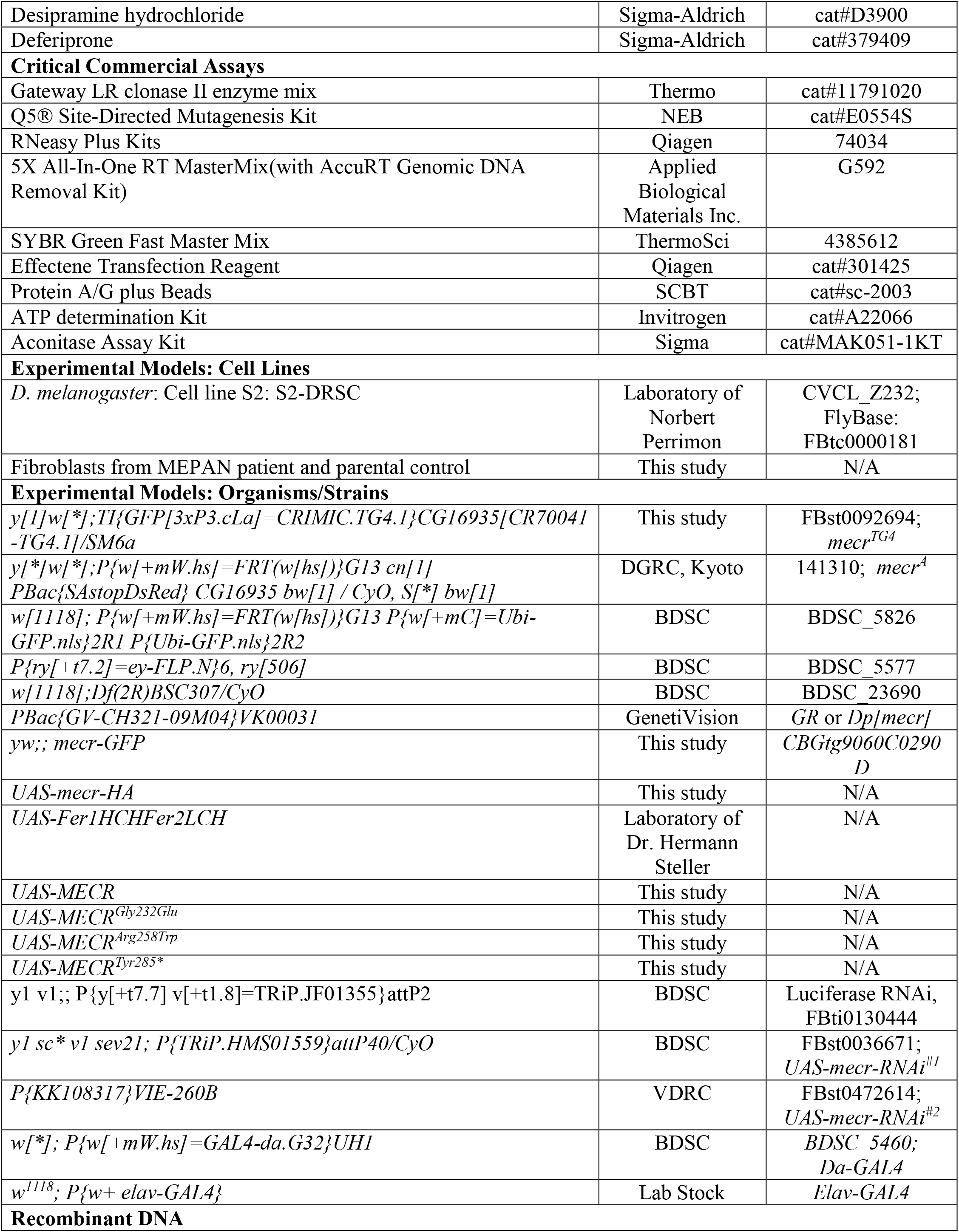
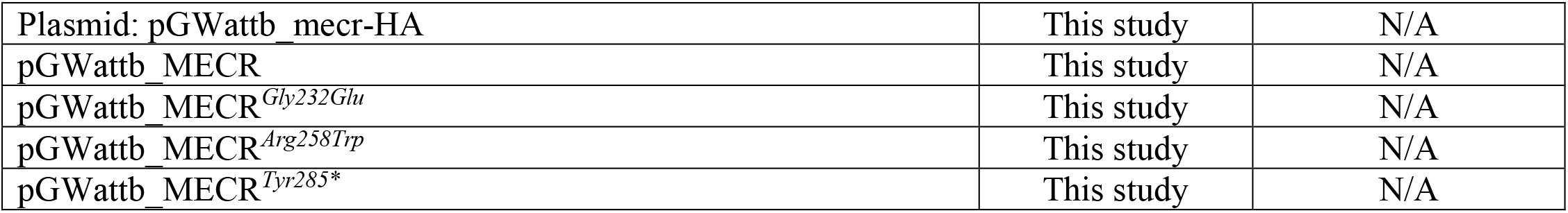

### Consortia

Undiagnosed Disease Network Consortia Member List: Mercedes E. Alejandro, Mahshid S. Azamian, Carlos A. Bacino, Ashok Balasubramanyam, Lindsay C. Burrage, Hsiao-Tuan Chao, Gary D. Clark, William J. Craigen, Hongzheng Dai, Shweta U. Dhar, Lisa T. Emrick, Alica M. Goldman, Neil A. Hanchard, Fariha Jamal, Lefkothea Karaviti, Seema R. Lalani, Brendan H. Lee, Richard A. Lewis, Ronit Marom, Paolo M. Moretti, David R. Murdock, Sarah K. Nicholas, James P. Orengo, Jennifer E. Posey, Lorraine Potocki, Jill A. Rosenfeld, Susan L. Samson, Daryl A. Scott, Alyssa A. Tran, Tiphanie P. Vogel, Michael F. Wangler, Shinya Yamamoto, Christine M. Eng, Pengfei Liu, Patricia A. Ward, Edward Behrens, Matthew Deardorff, Marni Falk, Kelly Hassey, Kathleen Sullivan, Adeline Vanderver, David B. Goldstein, Heidi Cope, Allyn McConkie-Rosell, Kelly Schoch, Vandana Shashi, Edward C. Smith, Rebecca C. Spillmann, Jennifer A. Sullivan, Queenie K.-G. Tan, Nicole M. Walley, Pankaj B. Agrawal, Alan H. Beggs, Gerard T. Berry, Lauren C. Briere, Laurel A. Cobban, Matthew Coggins, Cynthia M. Cooper, Elizabeth L. Fieg, Frances High, Ingrid A. Holm, Susan Korrick, Joel B. Krier, Sharyn A. Lincoln, Joseph Loscalzo, Richard L. Maas, Calum A. MacRae, J. Carl Pallais, Deepak A. Rao, Lance H. Rodan, Edwin K. Silverman, Joan M. Stoler, David A. Sweetser, Melissa Walker, Chris A. Walsh, Cecilia Esteves, Emily G. Kelley, Isaac S. Kohane, Kimberly LeBlanc, Alexa T. McCray, Anna Nagy, Surendra Dasari, Brendan C. Lanpher, Ian R. Lanza, Eva Morava, Devin Oglesbee, Guney Bademci, Deborah Barbouth, Stephanie Bivona, Olveen Carrasquillo,Ta Chen Peter Chang, Irman Forghani, Alana Grajewski, Rosario Isasi, Byron Lam, Roy Levitt, Xue Zhong Liu, Jacob McCauley, Ralph Sacco, Mario Saporta, Judy Schaechter,Mustafa Tekin, Fred Telischi, Willa Thorson, Stephan Zuchner, Heather A. Colley, Jyoti G. Dayal, David J. Eckstein, Laurie C. Findley, Donna M. Krasnewich, Laura A. Mamounas, Teri A. Manolio, John J. Mulvihill, Grace L. LaMoure, Madison P. Goldrich, Tiina K. Urv, Argenia L. Doss, Maria T. Acosta, Carsten Bonnenmann, Precilla D’Souza, David D. Draper, Carlos Ferreira, Rena A. Godfrey, Catherine A. Groden, Ellen F. Macnamara, Valerie V. Maduro, Thomas C. Markello, Avi Nath, Donna Novacic, Barbara N. Pusey, Camilo Toro, Colleen E. Wahl, Eva Baker, Elizabeth A. Burke, David R. Adams, William A. Gahl, May Christine V. Malicdan, Cynthia J. Tifft, Lynne A. Wolfe, John Yang, Bradley Power, Bernadette Gochuico, Laryssa Huryn, Lea Latham, Joie Davis, Deborah Mosbrook-Davis, Francis Rossignol, Ben Solomon, John MacDowall, Audrey Thurm, Wadih Zein, Muhammad Yousef, Margaret Adam, Laura Amendola, Michael Bamshad, Anita Beck, Jimmy Bennett, Beverly Berg-Rood, Elizabeth Blue, Brenna Boyd, Peter Byers,Sirisak Chanprasert, Michael Cunningham, Katrina Dipple, Daniel Doherty, Dawn Earl, Ian Glass, Katie Golden-Grant, Sihoun Hahn, Anne Hing, Fuki M. Hisama, Martha Horike-Pyne, Gail P. Jarvik, Jeffrey Jarvik, Suman Jayadev, Christina Lam, Kenneth Maravilla, Heather Mefford, J. Lawrence Merritt, Ghayda Mirzaa,Deborah Nickerson, Wendy Raskind, Natalie Rosenwasser, C. Ron Scott, Angela Sun, Virginia Sybert, Stephanie Wallace, Mark Wener, Tara Wenger, Euan A. Ashley, Gill Bejerano, Jonathan A. Bernstein, Devon Bonner, Terra R. Coakley, Liliana Fernandez, Paul G. Fisher, Laure Fresard, Jason Hom, Yong Huang, Jennefer N. Kohler, Elijah Kravets, Marta M. Majcherska, Beth A. Martin, Shruti Marwaha, Colleen E. McCormack, Archana N. Raja, Chloe M. Reuter, Maura Ruzhnikov, Jacinda B. Sampson, Kevin S. Smith, Shirley Sutton, Holly K. Tabor, Brianna M. Tucker, Matthew T. Wheeler, Diane B. Zastrow, Chunli Zhao, William E. Byrd, Andrew B. Crouse, Matthew Might, Mariko Nakano-Okuno, Jordan Whitlock, Gabrielle Brown, Manish J. Butte, Esteban C. Dell’Angelica, Naghmeh Dorrani, Emilie D. Douine, Brent L. Fogel, Irma Gutierrez, Alden Huang, Deborah Krakow, Hane Lee, Sandra K. Loo, Bryan C. Mak, Martin G. Martin, Julian A. Martínez-Agosto, Elisabeth McGee, Stanley F. Nelson, Shirley Nieves-Rodriguez, Christina G. S. Palmer, Jeanette C. Papp, Neil H. Parker, Genecee Renteria, Rebecca H. Signer, Janet S. Sinsheimer, Jijun Wan, Lee-kai Wang, Katherine Wesseling Perry, Jeremy D. Woods, Justin Alvey, Ashley Andrews, Jim Bale, John Bohnsack, Lorenzo Botto, John Carey, Laura Pace, Nicola Longo, Gabor Marth, Paolo Moretti, Aaron Quinlan, Matt Velinder, Dave Viskochi, Pinar Bayrak-Toydemir, Rong Mao, Monte Westerfield, Anna Bican, Elly Brokamp, Laura Duncan, Rizwan Hamid, Jennifer Kennedy, Mary Kozuira, John H. Newman, John A. PhillipsIII, Lynette Rives, Amy K. Robertson, Emily Solem, Joy D. Cogan, F. Sessions Cole, Nichole Hayes, Dana Kiley, Kathy Sisco, Jennifer Wambach, Daniel Wegner, Dustin Baldridge, Stephen Pak, Timothy Schedl, Jimann Shin, and Lilianna Solnica-Krezel.

## Extended Data

**Extended Data Figure 1:**
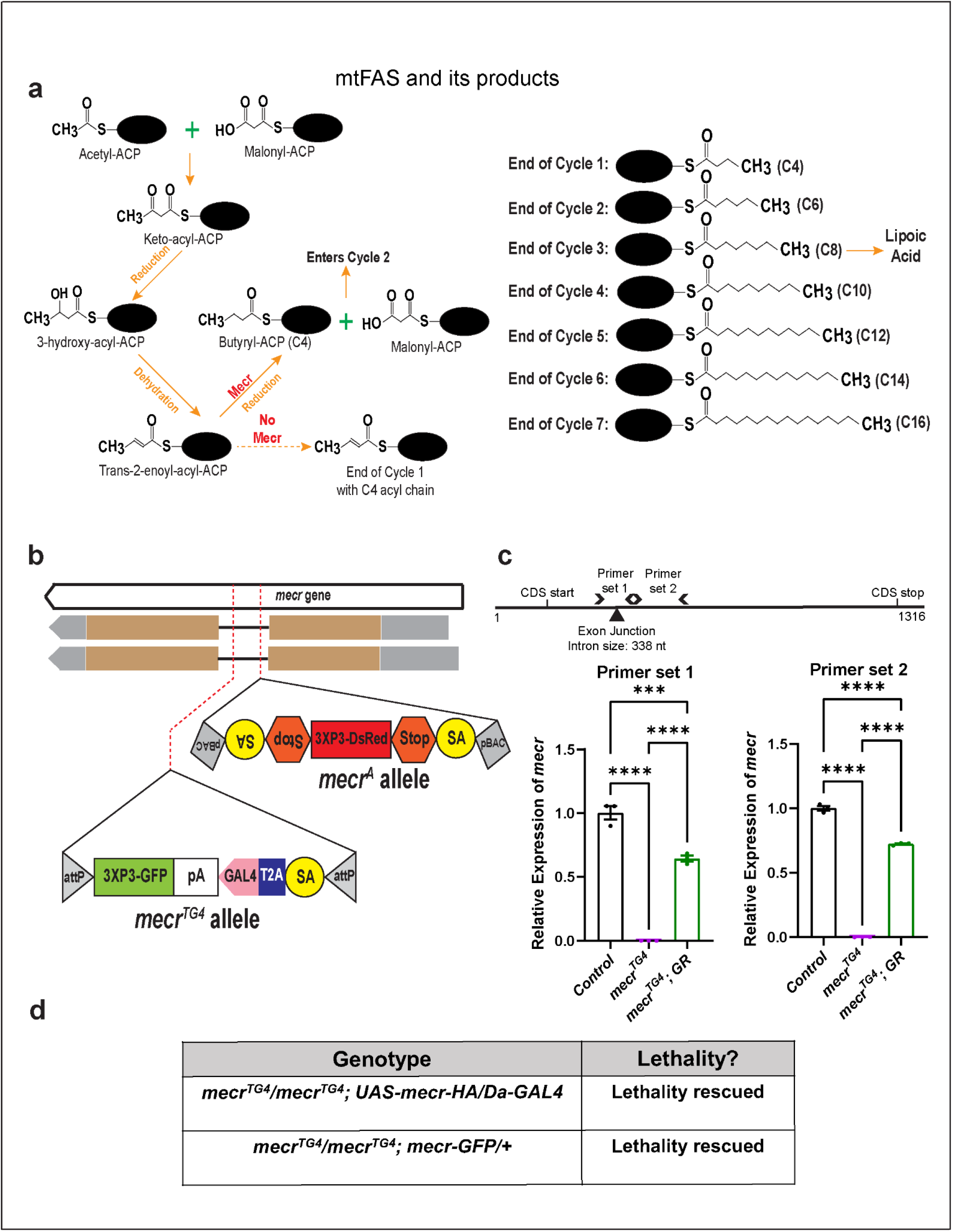
Mitochondrial fatty acid synthesis pathway and generation of *mecr* mutants. (a) Mitochondrial fatty acid synthesis pathway, its products and consequences upon loss of *mecr*. Briefly, an acetyl and malonyl moiety are condensed to make a four carbon long keto-acyl species, which remains attached to the mitochondrial Acyl Carrier Protein (mtACP), a central protein that holds the growing acyl chain during the fatty acid synthesis within the mitochondria. Subsequently, this four-carbon long keto-acyl ACP undergoes a reduction-dehydration-reduction cycle to produce a butyryl-ACP (C4). C4 enters into the cycle, and two carbons from the malonyl moiety are attached to the butyryl species to make an acyl chain of six carbon length. This cycle continues until the carbon length of the growing acyl chain reaches up to 16-18 carbon. (b) Schema showing the generation strategy of *mecr* mutants used in this study. (c) Schema showing the Primers used for quantitative real-time PCR and the graphs showing the relative levels of *mecr* transcripts in *mecr^TG4^*mutants. (d) Effects of a transgene containing *mecr-GFP* (Fosmid clone, CBGtg9060C0290D) ^36^ or of ubiquitous expression of HA-tagged fly *mecr* on *mecr^TG4^* homozygous mutants. One-way ANOVA followed by a Tukey’s post-hoc test was performed for the statistical analyses. Error bars represent SEM (****p < 0.0001).

**Extended Data Figure 2:**
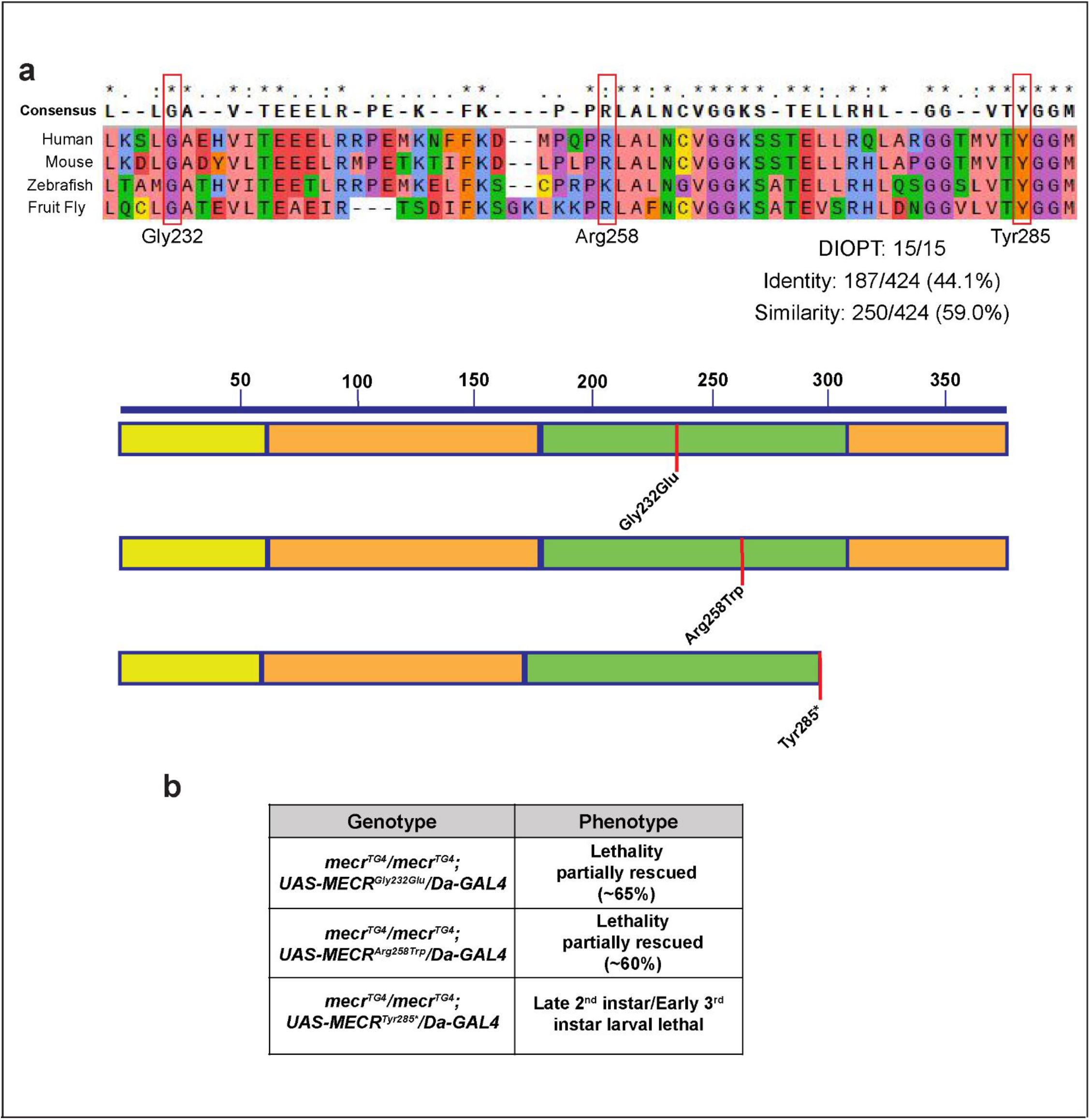
Lethality in homozygous *mecr* mutants is not rescued by expressing human MECR variants from patients. (a) Protein alignments of Mecr proteins. Red boxes indicate the amino acid that are mutated in MEPAN patients. Schema shows the relative position of the patient variants in MECR protein. (b) The effects of ubiquitous expression of human MECR mutations, which are observed in MEPAN patients, when driven by *Da-GAL4* in *mecr* mutants.

**Extended Data Figure 3:**
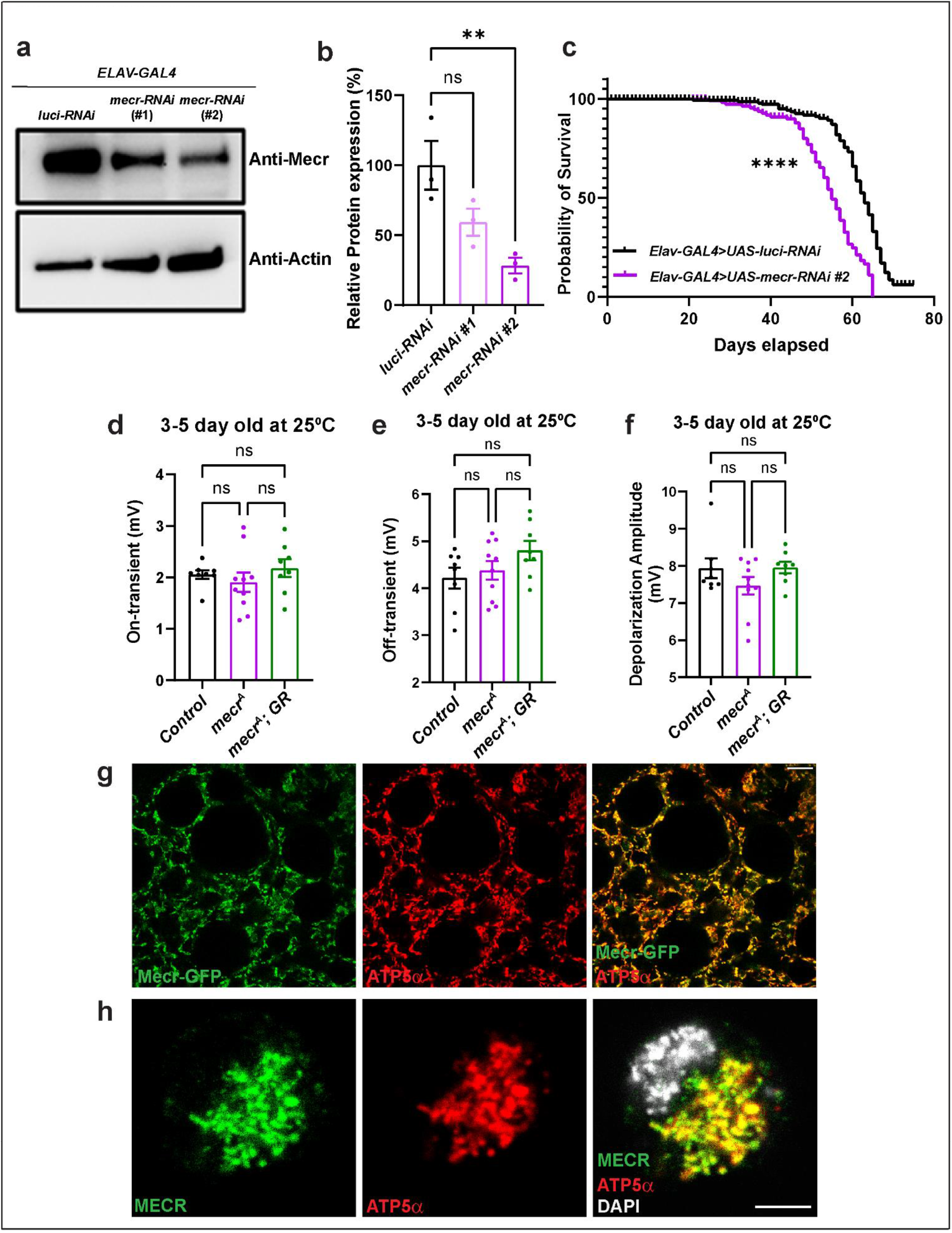
Loss of *mecr* causes a reduction in lifespan as well as climbing ability, and Fly Mecr and Human MECR proteins are localized to mitochondria. (a-b) Western blot and quantification showing relative levels of Mecr protein upon neuronal knockdown *(elav-GAL4)* with two different RNAi lines at 25°C. (c) Lifespan of flies with neuronal knockdown of *mecr*.n = 171 (*luci-RNAi*), n = 138 (*mecr-RNAi*) (d-f) Quantification of ERG traces from 3-5-day-old flies. n = 8 (*Control*), n = 10 (*mecr^A^*), n = 8 (*mecr^A^; GR*). For statistical analyses between two samples, two-tailed Student’s t-test is carried out. For statistical analyses among three samples one-way ANOVA followed by a Tukey’s post-hoc test is carried out. Error bars represent SEM (**p < 0.01; ****p < 0.0001). (g) Colocalization of Mecr-GFP and ATP5α in 3^rd^ instar larval fatbody tissue. Scale bar 10 µm. (h) Colocalization of human MECR and ATP5α in S2 cells. Scale bar 3.5 µm. Immunostaining was performed using an antibody against human MECR protein and an antibody against ATP5α.

**Extended Data Figure 4:**
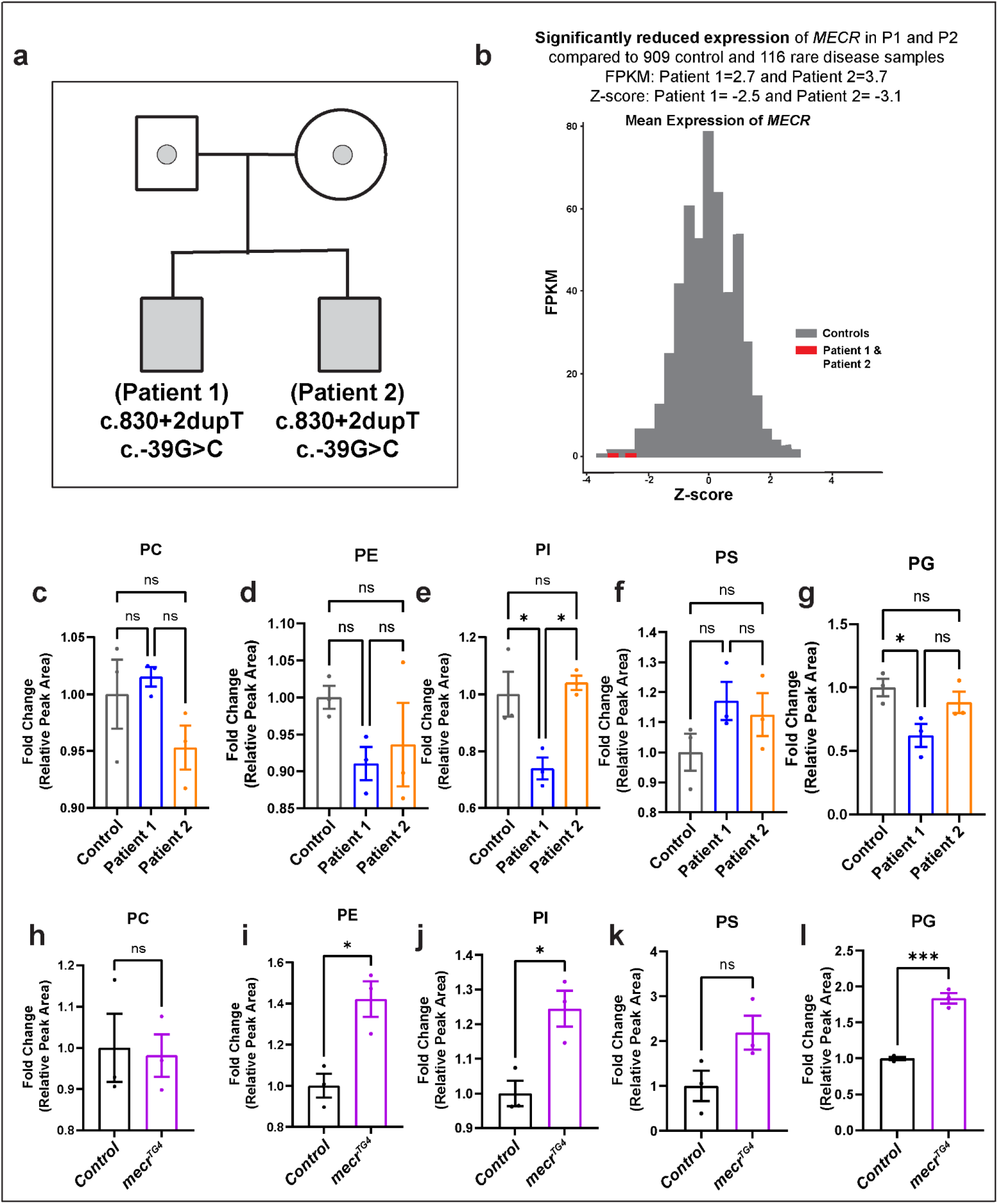
RNA levels of *MECR* are reduced in MEPAN patients and relative levels of phospholipids are differentially altered in patient-derived fibroblasts and *mecr* mutants. (a) Pedigree showing the two patients with MEPAN syndrome identified through UDN. (b) RNA-seq from blood showing reduced levels of *MECR* transcripts in both patients. (c-g) Relative phospholipids levels in MEPAN patient-derived fibroblasts compared to the parent-derived control fibroblasts: phosphatidylcholine (PC), phosphatidylethanolamine (PE), phosphatidylinositol (PI), phosphatidylserine (PS) and phosphatidylglycerol (PG). For statistical analyses one-way ANOVA followed by a Tukey’s post-hoc test are carried out. Error bars represent SEM (*p < 0.05) (h-l) Relative levels of different phospholipids in the *mecr^TG4^* larvae compared to control. For statistical two-tailed Student’s t-test are carried out. Error bars represent SEM (*p < 0.05; ***p < 0.001).

**Extended Data Figure 5:**
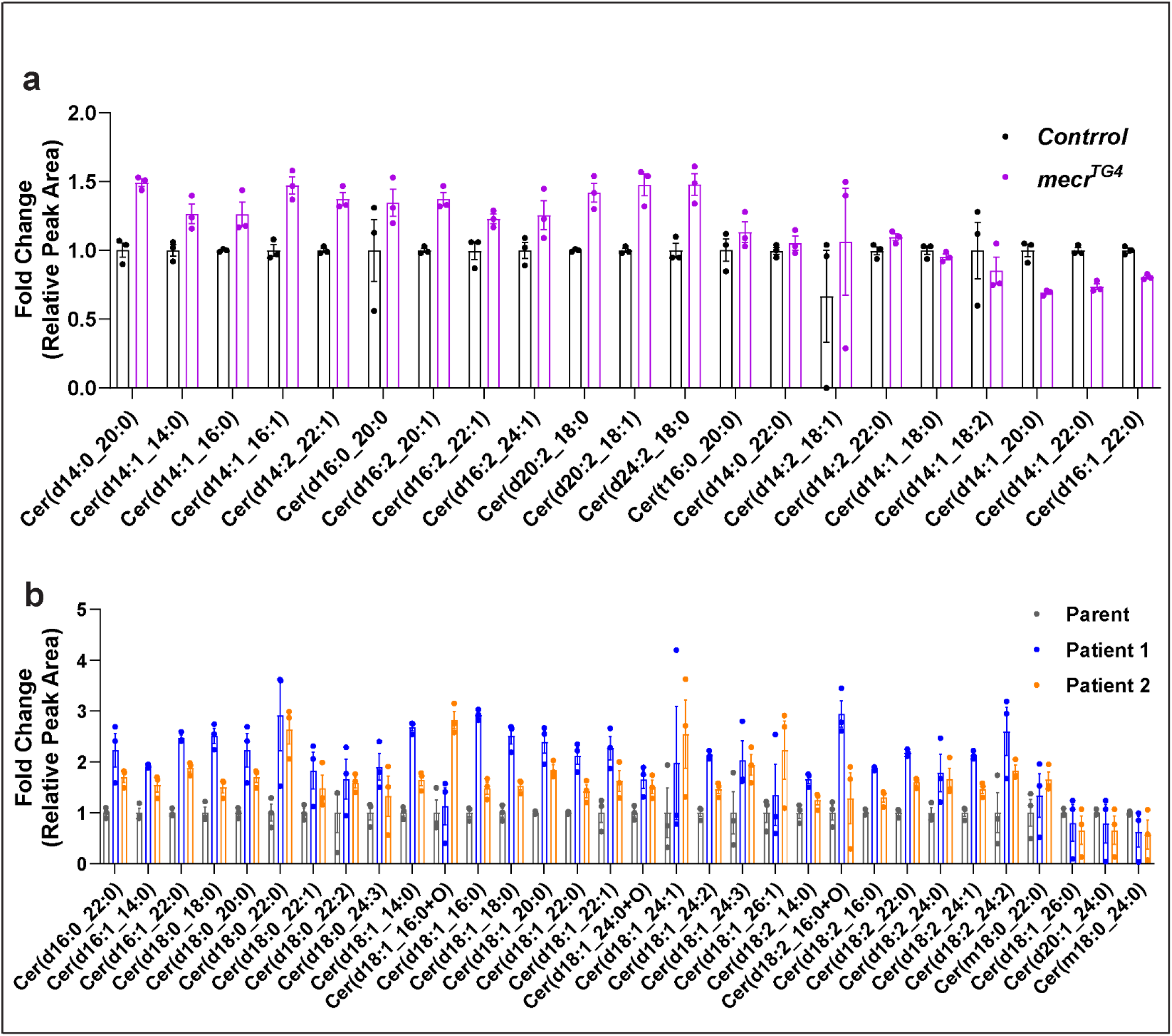
Altered ceramide levels in *mecr* mutants and human fibroblasts. (a) Graph showing the ceramide species that are elevated less than 1.5-fold or are not elevated in the *mecr^TG4^* fly mutant larvae. (b) Graph showing the ceramide species that are elevated less than 3-fold in both patient fibroblasts.

**Extended Data Figure 6:**
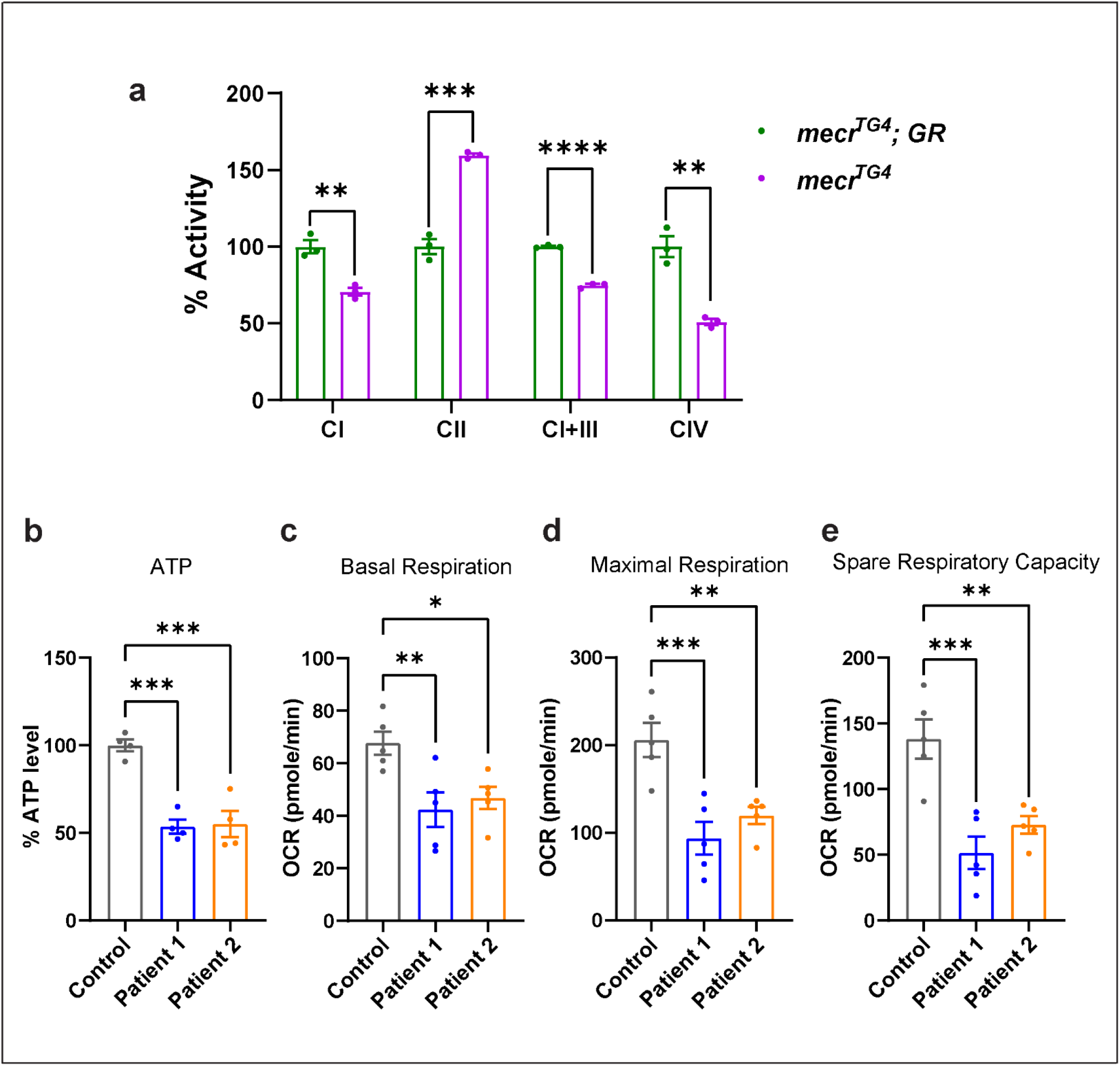
Loss of *mecr*/*MECR* leads to a respiratory deficit. (a) Relative activity of ETC complexes (CI-IV) in *mecr^TG4^* mutants and controls. *mecr^TG4^* mutant larvae display reduced activity of Complex-I, I+III, and IV and increased activity of Complex-II. (b) Relative levels of ATP in fibroblasts from patients and parental control. (c-e) Relative oxygen consumption rates in control and patient derived fibroblasts as measured by Seahorse analyses. (c) Basal respiration, (d) maximal respiration, and (e) spare respiratory capacity are reduced in the patient-derived fibroblasts compared to fibroblasts derived from parent control. For statistical analyses between two samples, two-tailed Student’s t test are carried out and for three samples one-way ANOVA followed by a Tukey’s post-hoc test are carried out. Error bars represent SEM (*p < 0.05; **p < 0.01; ***p < 0.001****p < 0.0001).

**Extended Data Figure 7:**
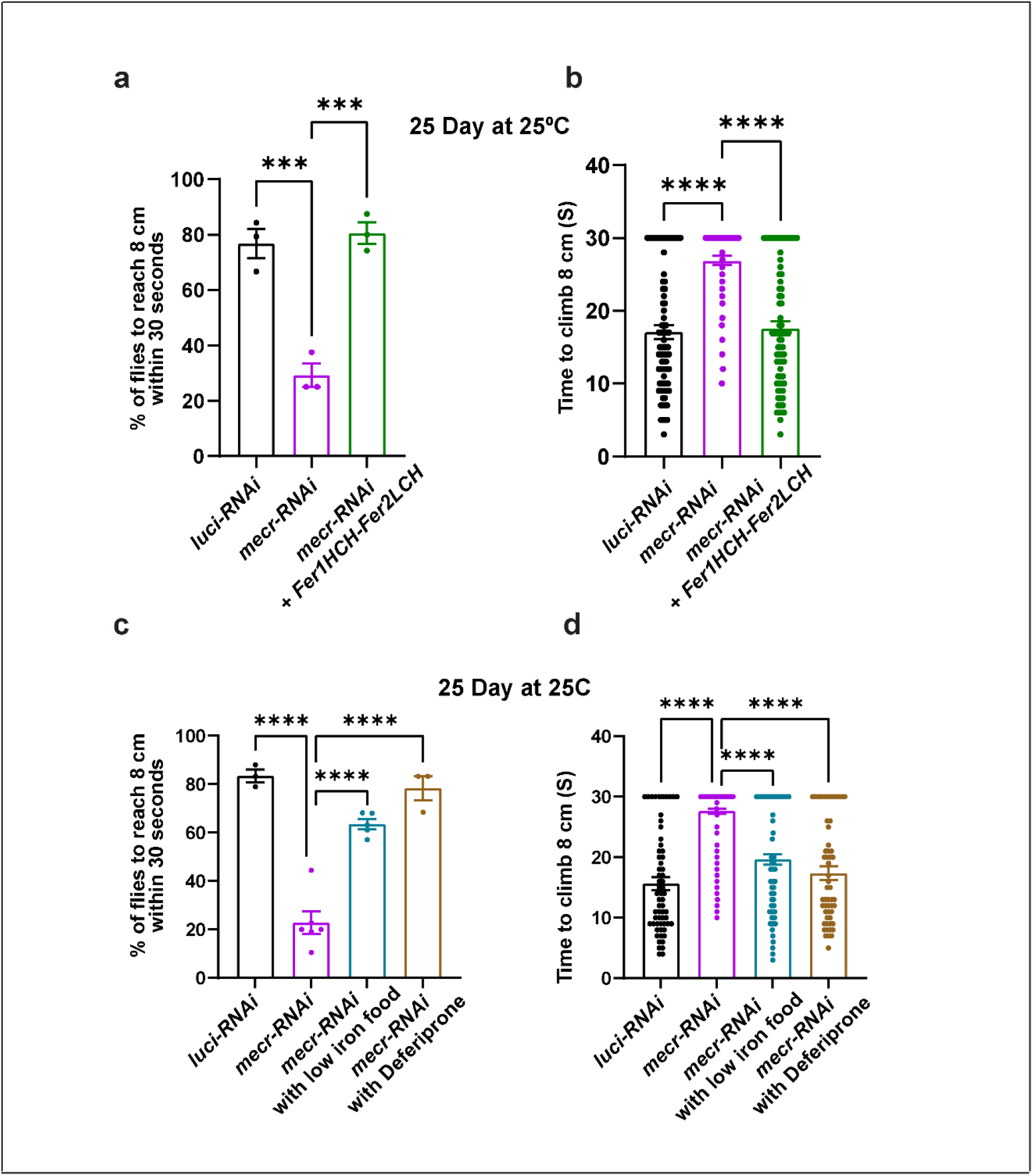
Reducing iron levels alleviates age-dependent locomotor defects in flies with neuronal knockdown of *mecr*. (a-b) Average percentage and climbing time of 25-day-old flies with neuronal (by *elav-GAL4*) *mecr-RNAi* and expressing *ferritins* to reach 8 cm. Total number of flies counted for three independent replicates were: n = 81 (*luci-RNAi*), n = 76 (*mecr-RNAi*), n = 74 (*mecr-RNAi* + *Fer1HCH-Fer2LCH*). (c-d) Average percentage and climbing time of 25-day-old flies upon neuronal (by *elav-GAL4*) knockdown of *mecr* treated with and without low iron food as well as Deferiprone. Total number of flies counted for three or more independent replicates were: n = 62 (*luci-RNAi*), n = 155 (*mecr-RNAi*), n = 105 (*mecr-RNAi* with low iron treatment), n = 55 (*mecr-RNAi* with low Deferiprone treatment). One-way ANOVA followed by a Tukey’s post-hoc test is carried out for statistical analyses. Error bars represent SEM (***p < 0.001; ****p < 0.0001).

**Extended Data Figure 8:**
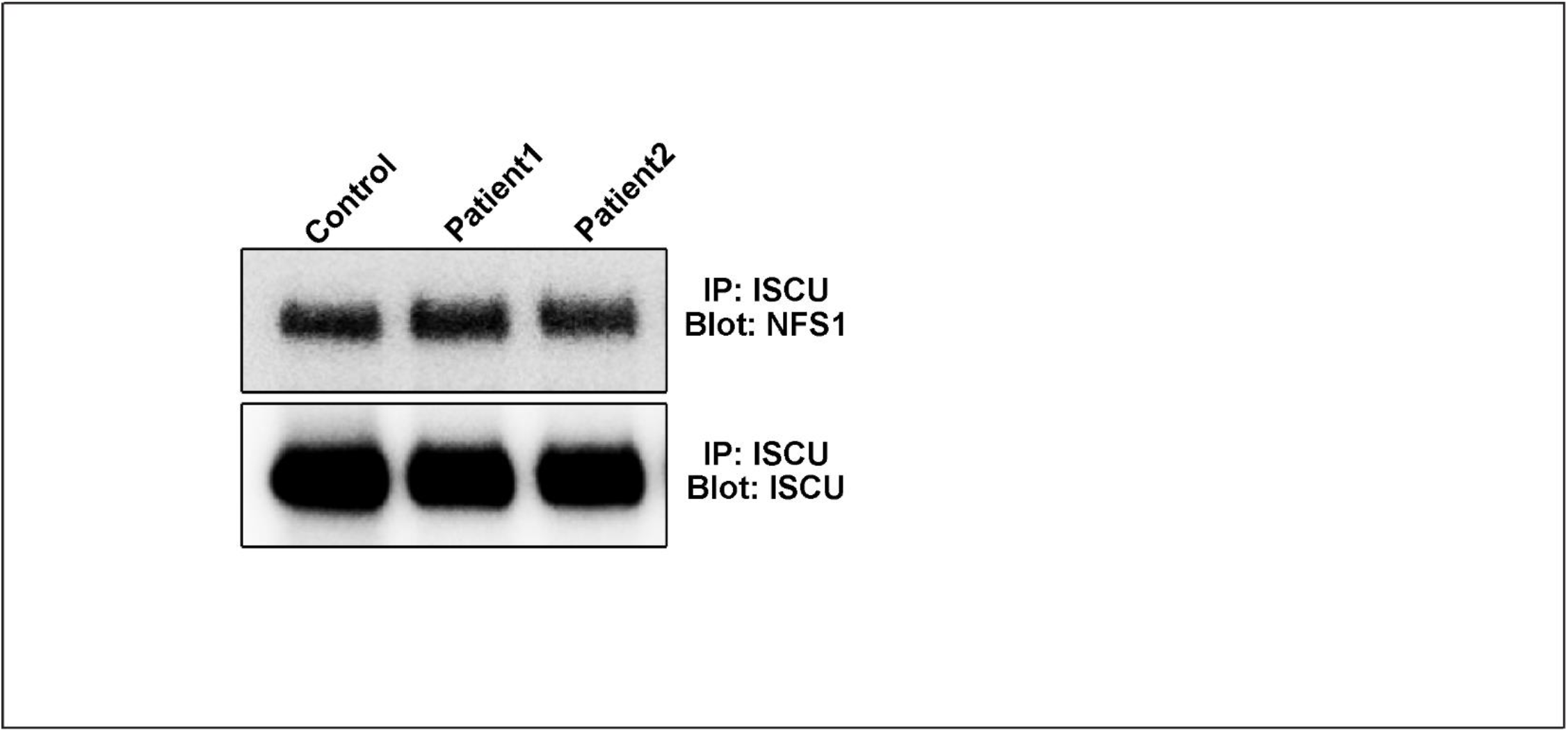
Co-IP shows the interaction between NFS1 and ISCU in the fibroblasts.

## References

1. Maier, T., Jenni, S. & Ban, N. Architecture of mammalian fatty acid synthase at 4.5 å resolution. Science (80-.). 311, 1258–1262 (2006).

2. Fhu, C. W. & Ali, A. Fatty Acid Synthase: An Emerging Target in Cancer. Molecules 25, 3935 (2020).

3. Miinalainen, I. J. et al. Characterization of 2-enoyl thioester reductase from mammals. An ortholog of YBR026p/MRF1’p of the yeast mitochondrial fatty acid synthesis type II. J. Biol. Chem. 278, 20154– 20161 (2003).

4. Nair, R. R. et al. Genetic modifications of Mecr reveal a role for mitochondrial 2-enoyl-CoA/ACP reductase in placental development in mice. Hum. Mol. Genet. 26, 2104–2117 (2017).

5. Venkatesan, R. et al. Insights into mitochondrial fatty acid synthesis from the structure of heterotetrameric 3-ketoacyl-ACP reductase/3R-hydroxyacyl-CoA dehydrogenase. Nat. Commun. 5, (2014).

6. Nowinski, S. M. et al. Mitochondrial fatty acid synthesis coordinates oxidative metabolism in mammalian mitochondria. Elife 9, 1–35 (2020).

7. Kursu, V. A. S. et al. Defects in mitochondrial fatty acid synthesis result in failure of multiple aspects of mitochondrial biogenesis in Saccharomyces cerevisiae. Mol. Microbiol. 90, 824–840 (2013).

8. Torkko, J. M. et al. Candida tropicalis Etr1p and Saccharomyces cerevisiae Ybr026p (Mrf1′p), 2-Enoyl Thioester Reductases Essential for Mitochondrial Respiratory Competence. Mol. Cell. Biol. 21, 6243– 6253 (2001).

9. Nair, R. R. et al. Impaired mitochondrial fatty acid synthesis leads to neurodegeneration in mice. J. Neurosci. 38, 9781–9800 (2018).

10. Heimer, G. et al. MECR Mutations Cause Childhood-Onset Dystonia and Optic Atrophy, a Mitochondrial Fatty Acid Synthesis Disorder. Am. J. Hum. Genet. 99, 1229–1244 (2016).

11. Gorukmez, O., Gorukmez, O. & Havalı, C. Novel MECR Mutation in Childhood-Onset Dystonia, Optic Atrophy, and Basal Ganglia Signal Abnormalities. Neuropediatrics 50, 336–337 (2019).

12. Liu, Z. et al. Whole exome sequencing identifies a novel homozygous MECR mutation in a Chinese patient with childhood-onset dystonia and basal ganglia abnormalities, without optic atrophy. Mitochondrion 57, 222–229 (2021).

13. Grassi, D. et al. Identification of a highly neurotoxic α-synuclein species inducing mitochondrial damage and mitophagy in Parkinson’s disease. Proc. Natl. Acad. Sci. U. S. A. 115, E2634–E2643 (2018).

14. Kastaniotis, A. J., Autio, K. J. & R. Nair, R. Mitochondrial Fatty Acids and Neurodegenerative Disorders. Neurosci. 27, 143–158 (2021).

15. Majmudar, J. D. et al. 4′-Phosphopantetheine and long acyl chain-dependent interactions are integral to human mitochondrial acyl carrier protein function. Medchemcomm 10, 209–220 (2019).

16. Zhang, L., Joshi, A. K., Hofmann, J., Schweizer, E. & Smith, S. Cloning, expression, and characterization of the human mitochondrial β-ketoacyl synthase: Complementation of the yeast cem1 knock-out strain. J. Biol. Chem. 280, 12422–12429 (2005).

17. Masud, A. J., Kastaniotis, A. J., Rahman, M. T., Autio, K. J. & Hiltunen, J. K. Mitochondrial acyl carrier protein (ACP) at the interface of metabolic state sensing and mitochondrial function. Biochim. Biophys. Acta - Mol. Cell Res. 1866, 118540 (2019).

18. Kastaniotis, A. J. et al. Mitochondrial fatty acid synthesis, fatty acids and mitochondrial physiology. Biochim. Biophys. Acta - Mol. Cell Biol. Lipids 1862, 39–48 (2017).

19. Solmonson, A. & DeBerardinis, R. J. Lipoic acid metabolism and mitochondrial redox regulation. J. Biol. Chem. 293, 7522–7530 (2018).

20. Hiltunen, J. K. et al. Mitochondrial fatty acid synthesis and respiration. Biochim. Biophys. Acta - Bioenerg. 1797, 1195–1202 (2010).

21. Schneider, R., Massow, M., Lisowsky, T. & Weiss, H. Different respiratory-defective phenotypes of Neurospora crassa and Saccharomyces cerevisiae after inactivation of the gene encoding the mitochondrial acyl carrier protein. Curr. Genet. 29, 10–17 (1995).

22. Guler, J. L., Kriegova, E., Smith, T. K., Lukeš, J. & Englund, P. T. Mitochondrial fatty acid synthesis is required for normal mitochondrial morphology and function in Trypanosoma brucei. Mol. Microbiol. 67, 1125–1142 (2008).

23. Clay, H. B. et al. Altering the mitochondrial fatty acid synthesis (mtFASII) pathway modulates cellular metabolic states and bioactive lipid profiles AS revealed by metabolomic profiling. PLoS One 11, 1–23 (2016).

24. Kanca, O. et al. An efficient CRISPR-based strategy to insert small and large fragments of DNA using short homology arms. Elife 8, 1–22 (2019).

25. Lee, P. T. et al. A gene-specific T2A-GAL4 library for drosophila. Elife 7, e35574 (2018).

26. Schuldiner, O. et al. piggyBac-Based Mosaic Screen Identifies a Postmitotic Function for Cohesin in Regulating Developmental Axon Pruning. Dev. Cell 14, 227–238 (2008).

27. Venken, K. J. T. et al. Versatile P[acman] BAC libraries for transgenesis studies in Drosophila melanogaster. Nat. Methods 6, 431–434 (2009).

28. Hu, Y. et al. An integrative approach to ortholog prediction for disease-focused and other functional studies. BMC Bioinformatics 12, 357 (2011).

29. Wang, J. et al. MARRVEL: Integration of Human and Model Organism Genetic Resources to Facilitate Functional Annotation of the Human Genome. Am. J. Hum. Genet. 100, 843–853 (2017).

30. Golic, K. G. & Lindquist, S. The FLP recombinase of yeast catalyzes site-specific recombination in the drosophila genome. Cell 59, 499–509 (1989).

31. Xu, T. & Rubin, G. M. Analysis of genetic mosaics in developing and adult Drosophila tissues. Development 117, 1223–1237 (1993).

32. Yamazoe, M. et al. A protein which binds preferentially to single-stranded core sequence of autonomously replicating sequence is essential for respiratory function in mitochondria of Saccharomyces cerevisiae. J. Biol. Chem. 269, 15244–15252 (1994).

33. Kim, D. G. et al. A novel cytosolic isoform of mitochondrial trans-2-enoyl-CoA reductase enhances peroxisome proliferator-activated receptor a activity. Endocrinol. Metab. 29, 185–194 (2014).

34. Masuda, N. et al. Nuclear receptor binding factor-1 (NRBF-1), a protein interacting with a wide spectrum of nuclear hormone receptors. Gene 221, 225–233 (1998).

35. Parl, A. et al. The mitochondrial fatty acid synthesis (mtFASII) pathway is capable of mediating nuclear-mitochondrial cross talk through the PPAR system of transcriptional activation. Biochem. Biophys. Res. Commun. 441, 418–424 (2013).

36. Sarov, M. et al. A genome-wide resource for the analysis of protein localisation in Drosophila. Elife 5, e12068 (2016).

37. Yoon, W. H. et al. Loss of Nardilysin, a Mitochondrial Co-chaperone for α-Ketoglutarate Dehydrogenase, Promotes mTORC1 Activation and Neurodegeneration. Neuron 93 115–131 (2017).

38. Baqri, R. M. et al. Disruption of mitochondrial DNA replication in Drosophila increases mitochondrial fast axonal transport in vivo. PLoS One 4, e7874 (2009).

39. Splinter, K. et al. Effect of Genetic Diagnosis on Patients with Previously Undiagnosed Disease. N. Engl. J. Med. 379, 2131–2139 (2018).

40. Ishibashi, Y., Kohyama-Koganeya, A. & Hirabayashi, Y. New insights on glucosylated lipids: Metabolism and functions. Biochim. Biophys. Acta - Mol. Cell Biol. Lipids 1831, 1475–1485 (2013).

41. Checa, A. et al. Hexosylceramides as intrathecal markers of worsening disability in multiple sclerosis. Mult. Scler. 21, 1271–1279 (2015).

42. Vos, M. et al. Ceramide accumulation induces mitophagy and impairs β-oxidation in PINK1 deficiency. Proc. Natl. Acad. Sci. U. S. A. 118, e2025347118 (2021).

43. Miyake, Y., Kozutsumi, Y., Nakamura, S., Fujita, T. & Kawasaki, T. Serine palmitoyltransferase is the primary target of a sphingosine-like immunosuppressant, ISP-1/myriocin. Biochemical and Biophysical Research Communications 211, 396–403 (1995).

44. Lin, G. et al. Phospholipase PLA2G6, a Parkinsonism-Associated Gene, Affects Vps26 and Vps35, Retromer Function, and Ceramide Levels, Similar to α-Synuclein Gain. Cell Metab. 28, 605–618.e6 (2018).

45. Jaiswal, M. et al. Impaired mitochondrial energy production causes light-induced photoreceptor degeneration independent of oxidative stress. PLoS Biol. 13, (2015).

46. Ozaki, M., Le, T. D. & Inoue, Y. H. Downregulating Mitochondrial DNA Polymerase γ in the Muscle Stimulated Autophagy, Apoptosis, and Muscle Aging-Related Phenotypes in Drosophila Adults. Biomolecules 12, 1105 (2022).

47. Larsen, S. B., Hanss, Z. & Krüger, R. The genetic architecture of mitochondrial dysfunction in Parkinson’s disease. Cell Tissue Res. 373, 21–37 (2018).

48. Zorova, L. D., et al. Mitochondrial membrane potential. Anal. Biochem. 552, 50–59 (2018).

49. Scaduto, R. C. & Grotyohann, L. W. Measurement of mitochondrial membrane potential using fluorescent rhodamine derivatives. Biophys. J. 76, 469–477 (1999).

50. Donti, T. R. et al. Screen for abnormal mitochondrial phenotypes in mouse embryonic stem cells identifies a model for succinyl-CoA ligase deficiency and mtDNA depletion. DMM Dis. Model. Mech. 7, 271–280 (2014).

51. Ji, R., et al. Increased de novo ceramide synthesis and accumulation in failing myocardium. JCI insight 2, e82922 (2017).

52. Srivastava, S. et al. SPTSSA Variants Alter Sphingolipid Synthesis and Cause a Complex Hereditary Spastic Paraplegia. Brain (Accepted Manuscript) (2022).

53. Chen, K. et al. Loss of frataxin activates the iron/sphingolipid/PDK1/Mef2 pathway in mammals. Elife 5, 1–14 (2016).

54. Chen, K. et al. Loss of frataxin induces iron toxicity, sphingolipid synthesis, and Pdk1/Mef2 activation, leading to neurodegeneration. Elife 5, 1–24 (2016).

55. Lee, Y. J. et al. Sphingolipid signaling mediates iron toxicity. Cell Metab. 16, 90–96 (2012).

56. Maio, N. & Rouault, T. A. Outlining the Complex Pathway of Mammalian Fe-S Cluster Biogenesis. Trends Biochem. Sci. 45, 411–426 (2020).

57. Meguro, R. et al. Nonheme-iron histochemistry for light end electorn microscopy: a historical, theoretical and technical review. Archives of Histology and Cytology 70, 1–19 (2007).

58. Galaris, D., Barbouti, A. & Pantopoulos, K. Iron homeostasis and oxidative stress: An intimate relationship. Biochim. Biophys. Acta - Mol. Cell Res. 1866, 118535 (2019).

59. Dixon, S. J. & Stockwell, B. R. The role of iron and reactive oxygen species in cell death. Nat. Chem. Biol. 10, 9–17 (2014).

60. Houglum, K., Filip, M., Witztum, J. L. & Chojkier, M. Malondialdehyde and 4-hydroxynonenal protein adducts in plasma and liver of rats with iron overload. J. Clin. Invest. 86, 1991–1998 (1990).

61. Csala, M. et al. On the role of 4-hydroxynonenal in health and disease. Biochim. Biophys. Acta - Mol. Basis Dis. 1852, 826–838 (2015).

62. Kruman, I., Bruce-Keller, A. J., Bredesen, D., Waeg, G. & Mattson, M. P. Evidence that 4- hydroxynonenal mediates oxidative stress-induced neuronal apoptosis. J. Neurosci. 17, 5089–5100 (1997).

63. Chung, H. lok, et al. Loss-or Gain-of-Function Mutations in ACOX1 Cause Axonal Loss via Different Mechanisms. Neuron 106, 589–606.e6 (2020).

64. Knovich, M. A., Storey, J. A., Coffman, L. G., Torti, S. V. & Torti, F. M. Ferritin for the clinician. Blood Rev. 23, 95–104 (2009).

65. Missirlis, F. et al. Homeostatic mechanisms for iron storage revealed by genetic manipulations and live imaging of Drosophila ferritin. Genetics 177, 89–100 (2007).

66. Soriano, S. et al. Deferiprone and idebenone rescue frataxin depletion phenotypes in a Drosophila model of Friedreich’s ataxia. Gene 521, 274–281 (2013).

67. Elincx-Benizri, S. et al. Clinical Experience with Deferiprone Treatment for Friedreich Ataxia. J. Child Neurol. 31, 1036–1040 (2016).

68. Vanlander, A. V. & Van Coster, R. Clinical and genetic aspects of defects in the mitochondrial iron– sulfur cluster synthesis pathway. J. Biol. Inorg. Chem. 23, 495–506 (2018).

69. Isaya, G. Mitochondrial iron-sulfur cluster dysfunction in neurodegenerative disease. Front. Pharmacol. 5, 1–7 (2014).

70. Van Vranken, J. G. et al. The mitochondrial acyl carrier protein (ACP) coordinates mitochondrial fatty acid synthesis with iron sulfur cluster biogenesis. Elife 5, 1–11 (2016).

71. Cory, S. A. et al. Structure of human Fe-S assembly subcomplex reveals unexpected cysteine desulfurase architecture and acyl-ACP-ISD11 interactions. Proc. Natl. Acad. Sci. U. S. A. 114, E5325–E5334 (2017).

72. Castro, L., Tórtora, V., Mansilla, S. & Radi, R. Aconitases: Non-redox Iron-Sulfur Proteins Sensitive to Reactive Species. Acc. Chem. Res. 52, 2609–2619 (2019).

73. Cheng, Z., Tsuda, M., Kishita, Y., Sato, Y. & Aigaki, T. Impaired energy metabolism in a Drosophila model of mitochondrial aconitase deficiency. Biochem. Biophys. Res. Commun. 433, 145–150 (2013).

74. Crooks, D. R. et al. Acute loss of iron–sulfur clusters results in metabolic reprogramming and generation of lipid droplets in mammalian cells. J. Biol. Chem. 293, 8297–8311 (2018).

75. Cho, Y. H., Kim, G. H. & Park, J. J. Mitochondrial aconitase 1 regulates age-related memory impairment via autophagy/mitophagy-mediated neural plasticity in middle-aged flies. Aging Cell 20, e13520 (2021).

76. Olona, A. et al. Sphingolipid metabolism during Toll-like receptor 4 (TLR4)-mediated macrophage activation. Br. J. Pharmacol. 178, 4575–4587 (2021).

77. Reginato, A. et al. The role of fatty acids in ceramide pathways and their influence on hypothalamic regulation of energy balance: a systematic review. Int. J. Mol. Sci. 22, 5357 (2021).

78. Nowinski, S. M., Van Vranken, J. G., Dove, K. K. & Rutter, J. Impact of Mitochondrial Fatty Acid Synthesis on Mitochondrial Biogenesis. Curr. Biol. 28, R1212–R1219 (2018).

79. Brody, S. & Mikolajczyk, S. Neurospora mitochondria contain an acyl-carrier protein. Eur. J. Biochem. 173, 353–359 (1988).

80. Stith, J. L., Velazquez, F. N. & Obeid, L. M. Advances in determining signaling mechanisms of ceramide and role in disease. J. Lipid Res. 60, 913–918 (2019).

81. Pinto, S. N., Silva, L. C., Futerman, A. H. & Prieto, M. Effect of ceramide structure on membrane biophysical properties: The role of acyl chain length and unsaturation. Biochim. Biophys. Acta - Biomembr. 1808, 2753–2760 (2011).

82. Alonso, A. & Goñi, F. M. The Physical Properties of Ceramides in Membranes. Annu. Rev. Biophys. 47, 633–654 (2018).

83. Rao, R. P. & Acharya, J. K. Sphingolipids and membrane biology as determined from genetic models. Prostaglandins Other Lipid Mediat. 85, 1–16 (2008).

84. Hernández-Corbacho, M. J., Salama, M. F., Canals, D., Senkal, C. E. & Obeid, L. M. Sphingolipids in mitochondria. Biochim. Biophys. Acta - Mol. Cell Biol. Lipids 1862, 56–68 (2017).

85. Kogot-Levin, A. & Saada, A. Ceramide and the mitochondrial respiratory chain. Biochimie 100, 88–94 (2014).

86. Alessenko, A. V. & Albi, E. Exploring Sphingolipid Implications in Neurodegeneration. Front. Neurol. 11, 437 (2020).

87. van Kruining, D. et al. Sphingolipids as prognostic biomarkers of neurodegeneration, neuroinflammation, and psychiatric diseases and their emerging role in lipidomic investigation methods. Adv. Drug Deliv. Rev. 159, 232–244 (2020).

88. Wang, D. et al. Skin fibroblast metabolomic profiling reveals that lipid dysfunction predicts the severity of Friedreich ’ s ataxia. J. Lipid Res. 63, 100255 (2022).

89. Anderson, P. R., Kirby, K., Hilliker, A. J. & Phillips, J. P. RNAi-mediated suppression of the mitochondrial iron chaperone, frataxin, in Drosophila. Hum. Mol. Genet. 14, 3397–3405 (2005).

90. Esposito, G. et al. Aconitase Causes Iron Toxicity in Drosophila pink1 Mutants. PLoS Genet. 9, (2013).

91. Ishihara-Paul, L. et al. PINK1 mutations and parkinsonism. Neurology 71, 896–902 (2008).

92. Custodia, A. et al. Ceramide Metabolism and Parkinson ’ s Disease — Therapeutic Targets. Biomolecules 11, 945 (2021).

93. Thomas, G. E. C. et al. Regional brain iron and gene expression provide insights into neurodegeneration in Parkinson’s disease. Brain 144, 1787–1798 (2021).

94. Foley, P. B., Hare, D. J. & Double, K. L. A brief history of brain iron accumulation in Parkinson disease and related disorders. J. Neural Transm. 129, 505–520 (2022).

95. Oakley, A. E. et al. Individual dopaminergic neurons show raised iron levels in Parkinson disease. Neurology 68, 1820–1825 (2007).

96. Wang, J. Y. et al. Meta-analysis of brain iron levels of Parkinson’s disease patients determined by postmortem and MRI measurements. Sci. Rep. 6, 36669 (2016).

97. Schweitzer, K. J. et al. Transcranial ultrasound in different monogenetic subtypes of Parkinson’s disease. J. Neurol. 254, 613–616 (2007).

98. Sawada, M. et al. P53 Regulates Ceramide Formation By Neutral Sphingomyelinase Through Reactive Oxygen Species in Human Glioma Cells. Oncogene 20, 1368–1378 (2001).

99. Li, X., Gulbins, E. & Zhang, Y. Oxidative stress triggers Ca 2+ -dependent lysosome trafficking and activation of acid sphingomyelinase. Cell. Physiol. Biochem. 30, 815–826 (2012).

100. Smith, A. R., Visioli, F., Frei, B. & Hagen, T. M. Lipoic acid significantly restores, in rats, the age-related decline in vasomotion. Br. J. Pharmacol. 153, 1615–1622 (2008).

101. Pavoine, C. & Pecker, F. Sphingomyelinases: Their regulation and roles in cardiovascular pathophysiology. Cardiovasc. Res. 82, 175–183 (2009).

102. Monette, J. S. et al. (R)-α-Lipoic acid treatment restores ceramide balance in aging rat cardiac mitochondria. Pharmacol. Res. 63, 23–29 (2011).

103. Schalinske, K. L. et al. The iron-sulfur cluster of iron regulatory protein 1 modulates the accessibility of RNA binding and phosphorylation sites. Biochemistry 36, 3950–3958 (1997).

104. Levi, S. & Rovida, E. Neuroferritinopathy: From ferritin structure modification to pathogenetic mechanism. Neurobiol. Dis. 81, 134–143 (2015).

105. Kumar, N., Rizek, P. & Jog, M. Neuroferritinopathy: Pathophysiology, presentation, differential diagnoses and management. Tremor and Other Hyperkinetic Movements 2016, 1–10 (2016).

106. Muhoberac, B. B. & Vidal, R. Iron, Ferritin, Hereditary Ferritinopathy, and Neurodegeneration. Front. Neurosci. 13, 1195 (2019).

107. Port, F., Chen, H. M., Lee, T. & Bullock, S. L. Optimized CRISPR/Cas tools for efficient germline and somatic genome engineering in Drosophila. Proc. Natl. Acad. Sci. U. S. A. 111, E2967–E2976 (2014).

108. Kanca, O. et al. An efficient CRISPR-based strategy to insert small and large fragments of DNA using short homology arms. Elife 8, e51539 (2019).

109. Dutta, D. et al. De novo mutations in TOMM70, a receptor of the mitochondrial import translocase, cause neurological impairment. Hum. Mol. Genet. 22, 1568–1579 (2020).

110. Marcogliese, P. C. et al. IRF2BPL Is Associated with Neurological Phenotypes. Am. J. Hum. Genet. 103, 245–260 (2018).

111. Marcogliese, P. C. et al. Loss of IRF2BPL impairs neuronal maintenance through excess Wnt signaling. Sci. Adv. 8, eabl5613 (2022).

112. Lee, J. W. et al. UPLC-QqQ/MS-Based Lipidomics Approach to Characterize Lipid Alterations in Inflammatory Macrophages. J. Proteome Res. 16, 1460–1469 (2017).

113. Radenkovic, S. et al. Expanding the clinical and metabolic phenotype of DPM2 deficient congenital disorders of glycosylation. Mol. Genet. Metab. 132, 27–37 (2021).

114. Byeon, S. K., Madugundu, A. K. & Pandey, A. Automated data-driven mass spectrometry for improved analysis of lipids with dual dissociation techniques. J. Mass Spectrom. Adv. Clin. Lab 22, 43–49 (2021).

115. Mitchell, C. J., Kim, M. S., Na, C. H. & Pandey, A. PyQuant: A versatile framework for analysis of quantitative mass spectrometry data. Mol. Cell. Proteomics 15, 2829–2838 (2016).

116. Frazier, A. E. & Thorburn, D. R. Biochemical Analyses of the Electron Transport Chain Complexes by Spectrophotometry. in Mitochondrial Disorders 837, 49–62 (2012).

117. Missirlis, F. et al. Characterization of mitochondrial ferritin in Drosophila. Proc. Natl. Acad. Sci. U. S. A. 103, 5893–5898 (2006).

118. Schober, F. A. et al. The one-carbon pool controls mitochondrial energy metabolism via complex i and iron-sulfur clusters. Sci. Adv. 7, (2021).

119. Martelli, F. et al. Low doses of the neonicotinoid insecticide imidacloprid induce ROS triggering neurological and metabolic impairments in Drosophila. Proc. Natl. Acad. Sci. U. S. A. 117, 25840–25850 (2020).

120. Dutta, D., Paul, M. S., Singh, A., Mutsuddi, M. & Mukherjee, A. Regulation of Notch Signaling by the Heterogeneous Nuclear Ribonucleoprotein Hrp48 and Deltex in Drosophila melanogaster. Genetics 206, 905–918 (2017).

## Extended Data References

Sarov, M. et al. A genome-wide resource for the analysis of protein localisation in Drosophila. Elife 5, e12068 (2016).

Heimer, G. et al. MECR Mutations Cause Childhood-Onset Dystonia and Optic Atrophy, a Mitochondrial Fatty Acid Synthesis Disorder. Am. J. Hum. Genet. 99, 1229–1244 (2016).

